# Type I partition-related proteins enhance conjugative transfer through transcriptional activation and *oriT* region binding

**DOI:** 10.1101/2025.01.22.634284

**Authors:** Kouhei Kishida, Koji Kudo, Ren Kumagai, Wenhao Deng, Leonardo Stari, Natsumi Ogawa-Kishida, Yoshiyuki Ohtsubo, Yuji Nagata, Masataka Tsuda

## Abstract

Plasmid partitioning and bacterial conjugation are critical processes ensuring plasmid maintenance and dissemination, respectively, within bacterial populations. Although traditionally regarded as distinct phenomena, these two processes are increasingly recognized as interconnected. While partitioning ensures plasmid inheritance during cell division, its potential influence on conjugative transfer remains poorly understood. A major impediment to understanding their interplay is that partition systems are often essential for plasmid stability, making it difficult to distinguish their direct effects on conjugation. In this study, we addressed this challenge using a mini-conjugative plasmid derived from the *Pseudomonas putida* NAH7 plasmid. This engineered plasmid, containing all conjugation-related genes, was cloned into an *E. coli*-compatible vector. Additionally, the *par* genes from NAH7 were expressed from a separate plasmid to investigate their roles in conjugative transfer. Our results revealed that the *par* gene cluster plays a significant role in enhancing the conjugative transfer of the mini-conjugative plasmid. Specifically, ParB, a centromere-binding protein, functions as a transcriptional activator of conjugation-related genes with binding *parS_NAH_* site. In contrast, ParR, a KorA homolog, was not found to enhance transcription directly but binds extensively to the *oriT* region. This binding probably facilitates the recruitment or stabilization of the relaxosome, thereby enhancing conjugation efficiency. Together, these findings unveil a previously unappreciated role for partition proteins in stimulating bacterial conjugation, providing new insights into how plasmids coordinate vertical and horizontal dissemination, highlighting that these processes can occur simultaneously within bacterial communities.

**IMPORTANCE:** Plasmid partition systems are classified into three types. While some systems have been reported to influence conjugative transfer, this study uncovers a novel mechanism utilized by a Type I system to enhance DNA transfer. Strikingly, repeat sequences perfectly matching the *parS_NAH_* site—bound by ParB to activate downstream conjugative transfer genes—were identified on both plasmids and chromosomes across diverse proteobacterial taxa. Furthermore, many of these repeat sequences were localized near genes involved in conjugative transfer and partitioning, suggesting the presence of a conserved regulatory mechanism mediated by these repeats. This study provides important insights into how plasmid partition systems coordinate both vertical and horizontal dissemination. Such knowledge is essential for understanding and mitigating the spread of antibiotic resistance and other plasmid-encoded traits, and it offers a foundation for developing strategies to manage plasmid-associated genetic exchange.

## INTRODUCTION

The propagation of plasmids through conjugative transfer is a socially significant phenomenon. It contributes to the emergence of antibiotic-resistant and pollutant-degrading bacteria (1), highlighting the need to understand the mechanisms that activate conjugative transfer. In Gram-negative bacteria, conjugation involves two main steps. First, the DNA transfer and replication (Dtr) system, in which relaxases serve as key enzymes, binds to and processes the cognate origin of transfer (*oriT*) DNA region, generating transfer-DNA (T-DNA) (1). The *oriT* DNA region is not only bound by relaxase but also by accessory proteins and transcriptional regulators that act on promoters located near the *oriT* region (2–4). Second, the conjugative proteins assemble into the type IV secretion system (T4SS) channel through which T-DNA is transferred to recipient cells (5, 6).

Low-copy-number plasmids are stably maintained by partition mechanisms that are responsible for their proper positioning within the cell during the cell division (7, 8). Plasmid partition systems (Par systems) generally comprise two essential proteins and partition sites (*par* sites) on plasmid DNA (7). One of these proteins, the centromere-binding protein (CBP), is a site-specific DNA-binding protein that recognizes the *par* sites. The other protein is an NTPase (ATPase or GTPase), which supplies the necessary energy to facilitate the movement of plasmid DNA within the cell. Par systems are classified into three types based on their NTPase properties: Type I (Walker-type ATPases), Type II (actin-like ATPases), and Type III (tubulin-like GTPases) (9, 10).

Par-related proteins (Par proteins) play roles in regulating conjugative transfer by repressing the transcription of conjugation-related genes and interacting with conjugation-associated proteins (11–14). Although Par proteins have been shown to enhance conjugative DNA transfer, the overall understanding of their roles in this process remains limited. One reason for this limitation is that disruption or overexpression of *par* genes inherently affects plasmid stability, making it difficult to isolate their direct effects on conjugation efficiency.

IncP-9 plasmid, which is primarily maintained in *Pseudomonas* species, often carries accessory genes, including antibiotic resistance genes and genes involved in the degradation of polycyclic aromatic hydrocarbons (15, 16). The IncP-9 naphthalene-catabolic plasmid NAH7 has been extensively studied for its conjugative transfer capabilities (17–22). It was originally isolated from *Pseudomonas putida* G7 (23), where it enables the host bacterium to utilize naphthalene as a source of carbon and energy. In practice, NAH7 can replicate and be maintained in both *Pseudomonas* and *Escherichia coli*, and it is capable of conjugative transfer not only to gamma-proteobacteria but also to alpha- and beta-proteobacteria, demonstrating a broad host range and further emphasizing its potential impact on microbial ecology and bioremediation (17). We have identified fundamental properties of NAH7 related to its conjugation system, including the *oriT*, its *nic* site, and the *relaxase* gene (17). pKKO3, a plasmid we constructed by cloning all the conjugation-related genes from NAH7 into the non-mobilizable shuttle vector pNIT6012 (24), has been demonstrated to be capable of conjugative transfer (18). Since pKKO3 does not contain the *par* genes from NAH7, it is suitable for investigating whether the *par* genes enhance conjugative transfer.

In this study, we demonstrated that the *par* gene cluster from the NAH7 Type I partition system, which includes the ParB as a CBP and ParR, the KorA homolog, contributes not only to plasmid stability but also to the activation of conjugative transfer. We found that ParB functions as a transcriptional activator for conjugation-related genes, likely through its binding to *parS_NAH_* sites. Although ParR did not exhibit transcriptional activation, it was observed to bind a broad region of the *oriT* region, suggesting that this association contributes to the enhancement of conjugative transfer. Together, our findings highlight a previously unappreciated role for Type I partition systems in stimulating conjugative transfer.

## Result

### Identification of plasmid partition-related genes and *par* sites on NAH7

To gain foundational insights into the Par system of NAH7, we conducted a bioinformatics analysis of its partition-related genes and associated *par* sites (Fig. 1A through C). Annotation of the NAH7 plasmid (Accession Number NC_007926.1) (25, 26) revealed a gene cluster comprising *parA*, *parB*, *parR*, and *parC* (Fig. 1A). ParA was predicted to be an ATPase containing a Walker A motif, characterized by the consensus sequence [A/G]XXXXG[S/T] (10), suggesting NAH7 utilizes a Type I partitioning system, and ParB was predicted as a CBP (Fig. S1). In Type I partitioning systems, the *par* sites recognized by CBPs typically contain inverted repeat (IR) sequences (10). Guided by this information, we searched for IR sequences in regions surrounding the *par* genes on NAH7 to locate potential *par* sites. This analysis identified an 18-bp IR sequence, 5′-TTTCTCGCATGCGAGAAA-3′ (designated *parS1_NAH_*), located at positions 82,127–82,143 (Fig. 1A and B).

**Figure 1.**
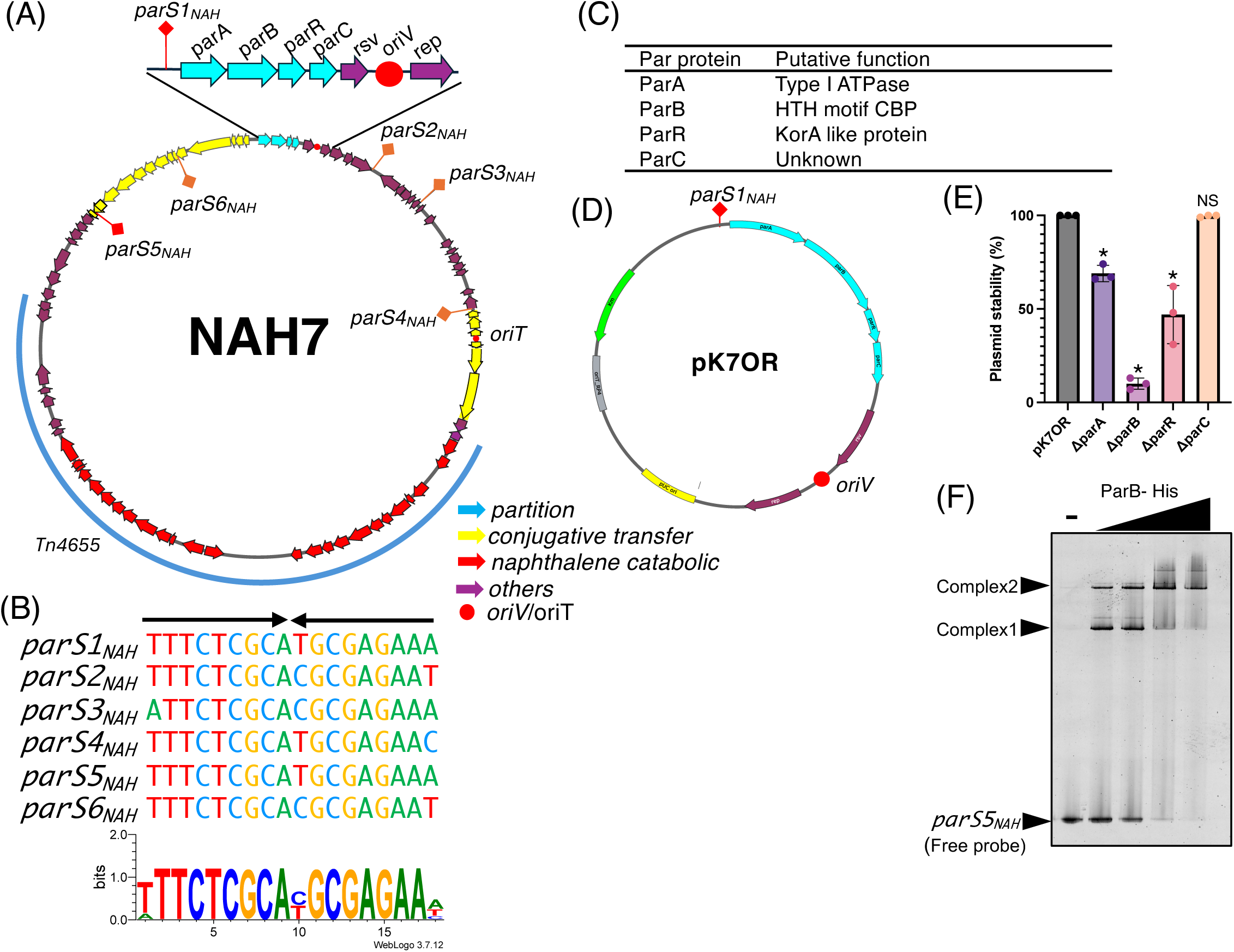
Overview of NAH7 plasmid partition system and characterization of *parS* sites and Par proteins. **(A)** Circular map of NAH7. Partitioning genes (*parA*, *parB*, *parR*, and *parC*) are indicated in cyan, conjugative transfer-related genes are indicated in yellow, naphthalene catabolic genes are shown in red, and other genes are shown in purple. The origin of replication *(oriV*) and origin of transfer (*oriT*) are marked with red circles. Six inverted repeat sequences, designated *parS1_NAH_* through *parS6_NAH_*, are distributed across the plasmid (marked in orange or red). **(B)** Alignment of the identified *parS_NAH_* sites. *parS_NAH_* sequences are shown with similarities highlighted, indicating conserved nucleotide patterns, *parS1_NAH_* and *parS5_NAH_* are perfect inverted repeat sequences, while the other *parS_NAH_* sites (*parS2_NAH_* through *parS4_NAH_*, *parS6_NAH_*) contain up to two mismatches. The sequence logo was generated using *parS1_NAH_* to *parS6_NAH_* with WebLogo (47). **(C)** Summary table listing the putative functions of the NAH7 partition proteins, based on predictions from InterPro analysis (Fig. S1). **(D)** Plasmid map of pK7OR, derived from pK18mob, highlighting key genetic elements. The plasmid features the *parS1_NAH_* site located upstream of the *parA*, *parB*, *parR*, and *parC* genes (cyan arrows). The *oriV* (red circle), *resolvase* and *rep* genes (purple arrows) together support plasmid replication, ensuring its maintenance in host cells. The Km gene (green arrow) provides kanamycin resistance for selection. **(E)** Plasmid stability assay for pK7OR, with deletions of individual *par* genes (Δ*parA*, Δ*parB*, Δ*parR*, Δ*parC*) conducted in *P. putida* KT2440. Stability was measured as the percentage of plasmid-harboring (Km^R^) colonies out of 100 colonies after 24 hours of growth in non-selective conditions The assay was performed in three independent biological replicates. The average values were plotted with individual data points, and standard deviations are represented as error bars. Statistical analyses were performed by ordinary one-way ANOVA with Dunnett’s multiple comparisons test, comparing each deletion mutant with pK7OR (∗: p < 0.005). "NS" denotes no significant difference. **(F)** Electrophoretic Mobility Shift Assay showing the binding of ParB-His protein to the *parS5_NAH_* DNA fragment. Each lane contains 23 nM *parS5_NAH_* DNA fragment, and the final concentrations of ParB-His were 0, 0.15, 0.3, 0.45, and 0.9 µM from the leftmost lane. Samples were electrophoresed on a 15% polyacrylamide gel at 200 V for 90 minutes using Tris-glycine buffer. Complex 1 and Complex 2 represent the major shifted bands, while *parS5_NAH_* represents the free probe.

Additionally, we discovered five other similar IR sequences (*parS2_NAH_* to *parS6_NAH_*), each differing by up to two nucleotides from *parS1_NAH_*, distributed across NAH7 and located in intergenic regions. Notably, these IR sequences were located in the IncP-9 backbone; none of them were found within the Tn*4655* transposable element, suggesting that these IR sequences are essential components of the plasmid backbone. The *parS5_NAH_* sequence, positioned upstream of the *<u>mpfK</u>* gene—an essential gene for conjugative transfer in IncP-9 plasmids—was identical to *parS1_NAH_*, while the other IRs exhibited partial conservation (Fig. 1B). Further analysis predicted that ParR contains a KorA domain (Fig. S1), known to be involved in transcriptional regulation. Although the specific function of ParC remains unknown, it is conserved among IncP-9 plasmids (Table S2).

To investigate the role of these *par* genes in maintaining the stability of NAH7, we performed plasmid stability assays using mini-NAH7 variants lacking each *par* gene in *P. putida* KT2440 (Fig. 1D). The mini-NAH7 plasmid (pK7OR) was engineered by cloning the region spanning from *parS1_NAH_* to the *rep* gene into an *E. coli* plasmid vector containing a kanamycin resistance marker (Fig. 1D). The mini-NAH7 plasmid exhibited stable inheritance, with 100% of the progeny retaining the plasmid after 24 hours of incubation. In contrast, deletions of *parA*, *parB*, and *parR* led to significantly reduced plasmid stability, whereas deletion of *parC* did not markedly affect stability. These results indicate that *parA*, *parB*, and *parR* are essential for plasmid stability, while *parC* is dispensable under the conditions tested.

To confirm that ParB functions as a CBP and that the *parS_NAH_* sequence acts as a *par* site on NAH7, we performed an Electrophoretic Mobility Shift Assay (EMSA) with a *parS_NAH_* DNA probe. Increasing concentrations of ParB resulted in the formation of two distinct DNA-protein complexes, designated Complex 1 and Complex 2. The presence of these complexes suggests that ParB binds to *parS_NAH_* in a multimeric form. These results support the conclusion that ParB is a CBP and that *parS_NAH_* functions as a *par* site essential for the partitioning system of NAH7.

### Enhanced conjugative transfer of pKKO3-2 by *par* genes

We found that the identified *parS_NAH_* sequences were not only located around the *par* genes but were also positioned close to genes involved in conjugative transfer (Fig. 1A). This distribution suggests that potential crosstalk between partitioning and conjugative transfer mechanisms in NAH7. Attempts to replace the *par* gene cluster in the NAH7 plasmid with a kanamycin resistance cassette yielded Km-resistant colonies. However, even after multiple rounds of single-colony isolation, every colony still harbored both wild-type NAH7 and the *par-* deleted plasmid (Fig. S2). Therefore, we employed a plasmid pKKO3-2 to investigate how *par* genes influence conjugative transfer (Fig. 2). pKKO3-2 carries all of the conjugation-related genes from NAH7 but lacks replication and partitioning genes, making it an ideal model for examining the direct effects of *par* genes on conjugative transfer. To assess the impact of *parABRC* on pKKO3-2 transfer between *E. coli* strains, we induced their expression from a pUC18 vector using IPTG. Under these conditions, the pKKO3-2 transfer frequency increased by more than 200-fold (Fig. 2B) without affecting donor cell growth. To identify which specific *par* genes were responsible for this enhancement (Fig. 2C), we expressed each gene individually. Expression of *parA*, *parB*, or *parR* significantly elevated transfer frequency, whereas *parC* did not. We further examined various two- and three-gene combinations (Fig. 2D and E), all of which produced higher transfer frequencies than any single gene alone. Notably, the *parA* and *parR* pair yielded the greatest increase among the two-gene sets, and among the three-gene combinations, *parARC* resulted in the highest transfer frequency. These findings indicate that *parA*, *parB*, and *parR* individually promote conjugative transfer, and that their combined expression can further intensify this effect.

**Fig. 2.**
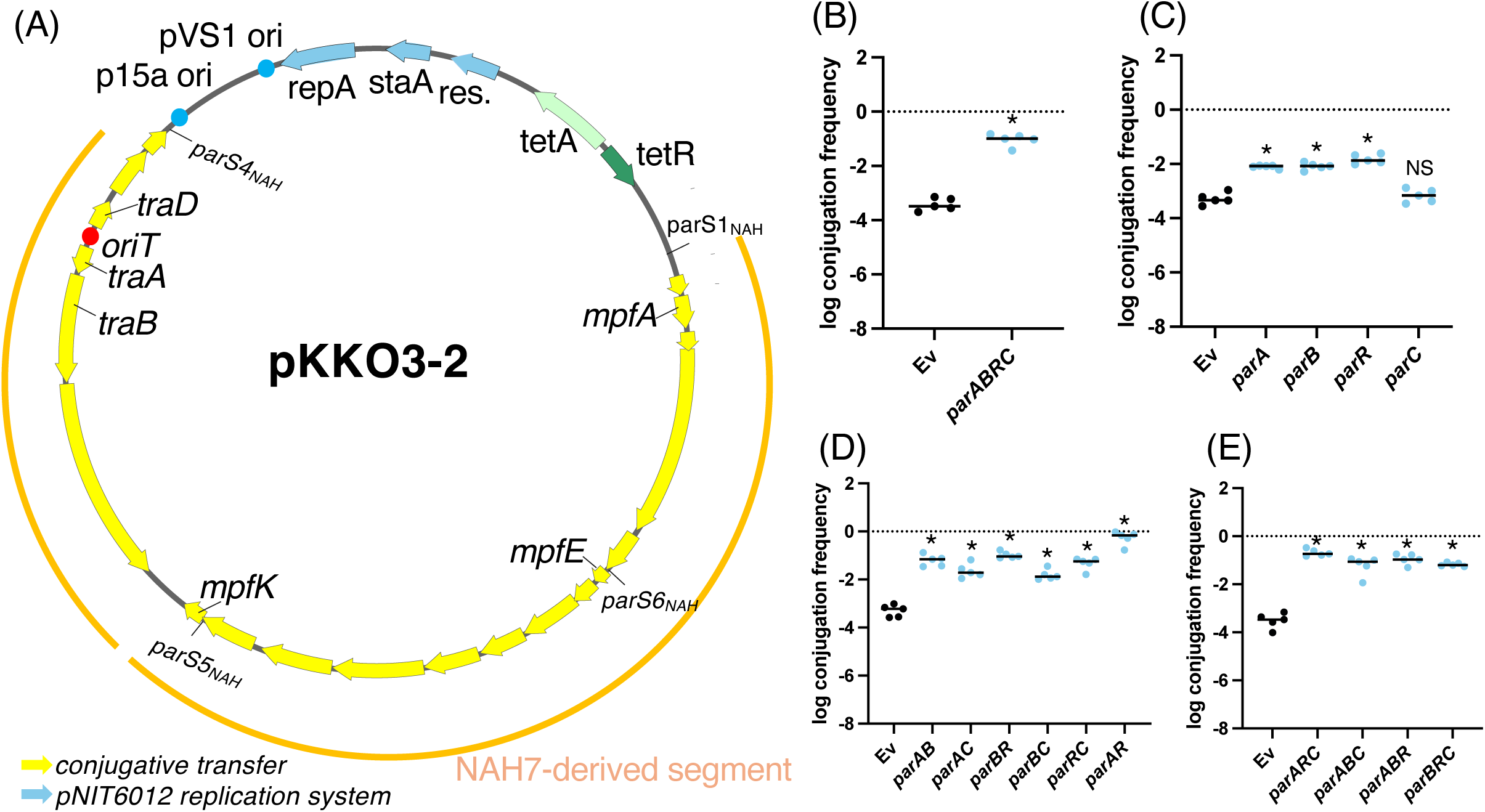
Effects of *par* genes on conjugative transfer of pKKO3-2. **(A)** Plasmid map of the pKKO3-2. The yellow segments represent genes involved in conjugative transfer derived from NAH7, and the blue arrows represent the pNIT6012 replication system. The locations of four *parS_NAH_* sites are shown. **(B–E)** Conjugative transfer frequency of pKKO3-2 with the expression of different *par* genes from a pUC18 vector. Panel (B) shows the effect of expressing all four *par* genes (*parA*, *parB*, *parR*, *parC*) together, while (C), (D), and (E) represent the effects of expressing different single *par* genes, as well as various combinations of two or three par genes. Conjugation frequency is shown on a log scale as the number of transconjugants per donor cell (Tc/D). Each experiment was performed at least five times independently. Error bars represent the standard deviation from biological replicates, with individual data points also plotted. Statistical analysis for Panel (B) was performed using an unpaired two-tailed t-test, while Panels (C), (D), and (E) were analyzed by ordinary one-way ANOVA followed by Dunnett’s multiple comparisons test relative to the strain harboring an empty vector (Ev). Asterisks (*, P < 0.0001) indicate significant differences compared to Ev. “NS” denotes no significant difference.

### Transcriptional analysis of conjugation-related genes in pKKO3-2 expressing individual *par* genes

Because several Par proteins are predicted to bind DNA, we hypothesized that they activate transcription of conjugation-related genes. To test this hypothesis, we used quantitative reverse transcription PCR (qRT-PCR) with *gyrA* as an internal standard to measure the transcription levels of several conjugation-related genes in *E. coli* strains harboring pKKO3-2, each expressing a single *par* gene. We quantified the relative expression of *mpfA*, *mpfE*, *mpfK*, *traD*, and *traB* (Fig. 3). Our results showed that T4SS-related genes (*mpfA*, *mpfE*, and *mpfK*) were upregulated in strains expressing *parA* or *parB* (Fig. 3A to C). In contrast, neither *parR* nor *parC* affected the transcription of any T4SS-related genes. We next examined *traD* and *traB*, which have promoters located on *oriT*. Here, an increase in transcription was observed only with *parA*, while the other *par* genes had no noticeable effect (Fig. 3D, E). *parA* increased the transcription of all conjugation-related genes tested. Interestingly, *parA* also elevated *tetA* transcription in the pNIT6012 segment (Fig. 3F), prompting us to investigate whether *parA* influences plasmid copy number. Indeed, only the *parA*-expressing strain showed an increase in pKKO3-2 copy number (Fig. 3G). To determine if this effect was specific to pKKO3-2, we assessed *parA*’s impact on pNIT6012 copy number, which also rose in the presence of *parA* (Fig. 3H). Although the mechanism underlying this copy number increase remains unclear, our findings suggest that *parA*-mediated transcriptional activation may be linked to higher plasmid copy number. Meanwhile These qRT-PCR data indicate that ParB likely acts as a transcriptional activator for genes downstream of *parS_NAH_* sequences (*mpfA*, *mpfE*, and *mpfK*), presumably by binding these sites.

**Fig. 3.**
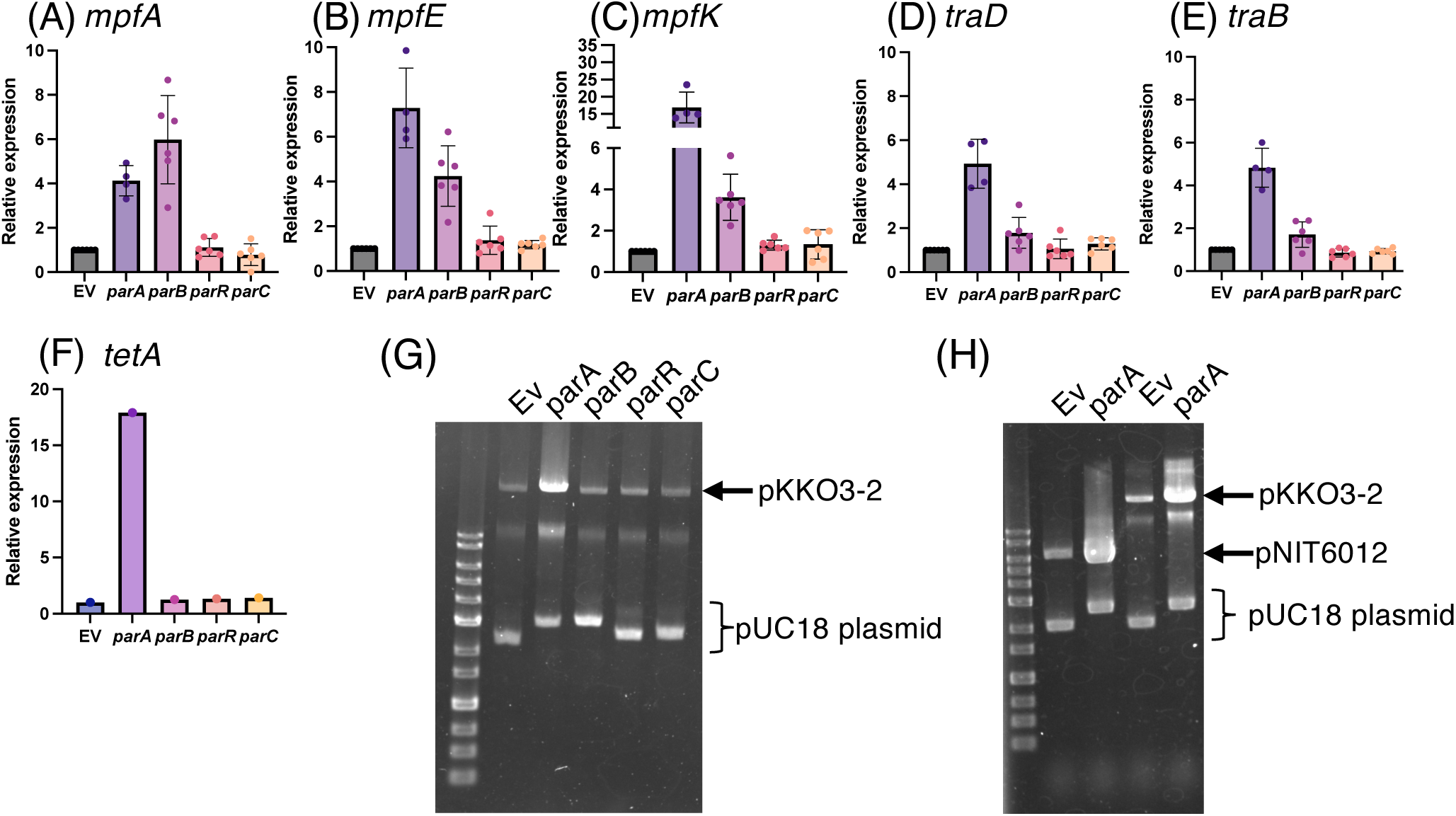
Relative expression levels of conjugative transfer genes in *E. coli* strains harboring the pKKO3-2 plasmid with individual *par* gene expression. **(A-F)** Gene expression levels were quantified using qRT-PCR with *gyrA* as the internal control. The graphs show the relative expression of (A) *mpfA*, (B) *mpfE*, (C) *mpfK*, (D) *traD*, (E) *traB* and (F) *tetA*. The relative expression values are normalized to the empty vector strain (EV), which is set as 1. The data represent the mean, with each dot corresponding to an independent biological replicate, with individual data points plotted. Error bars indicate standard deviation. **(G, H)** Agarose gel electrophoresis of plasmid isolation from *E. coli* strains. (G) shows plasmid profiles for *E. coli* carrying pKKO3-2 and pUC18 derivatives with different *par* gene constructs, while (H) shows profiles for *E. coli* harboring pNIT6012 and pUC18 derivatives. The specific constructs expressing *parA*, *parB*, *parR*, or *parC* are indicated above each lane.

### Distribution of *parS_NAH_* sites around conjugative transfer and partitioning genes in specific genera

Given that ParB binds to *parS_NAH_* and increases transcription levels downstream of these sites, we hypothesized that these sites might serve as regulatory hubs integrating vertical and horizontal plasmid dissemination. To test this hypothesis, we investigated the conservation and distribution of *parS_NAH_* sequences. We analyzed both plasmids and chromosomes containing at least one perfectly matching *parS1_NAH_* site sequence. As a result, we identified 120 plasmid sequences and 38 chromosomal sequences that contained a perfect *parS1_NAH_* match. For phylogenetic analysis, we constructed a tree using conserved genes from these representative sequences: *relaxase* genes were used for plasmid sequences, whereas 16S rRNA genes were used for chromosomal sequences, enabling classification and visualization by genus (Fig. 4A and B). Plasmid and chromosomal sequences were predominantly associated with genera such as *Pseudomonas*, *Xanthomonas*, *Burkholderia*, and *Ralstonia*. Interestingly, the analyzed *parS_NAH_* site was not only found in the plasmid of NAH7’s original host, *Pseudomonas*, but also in plasmids and chromosomes of several other proteobacterial species, highlighting its broader distribution and potential functional significance.

**Figure 4.**
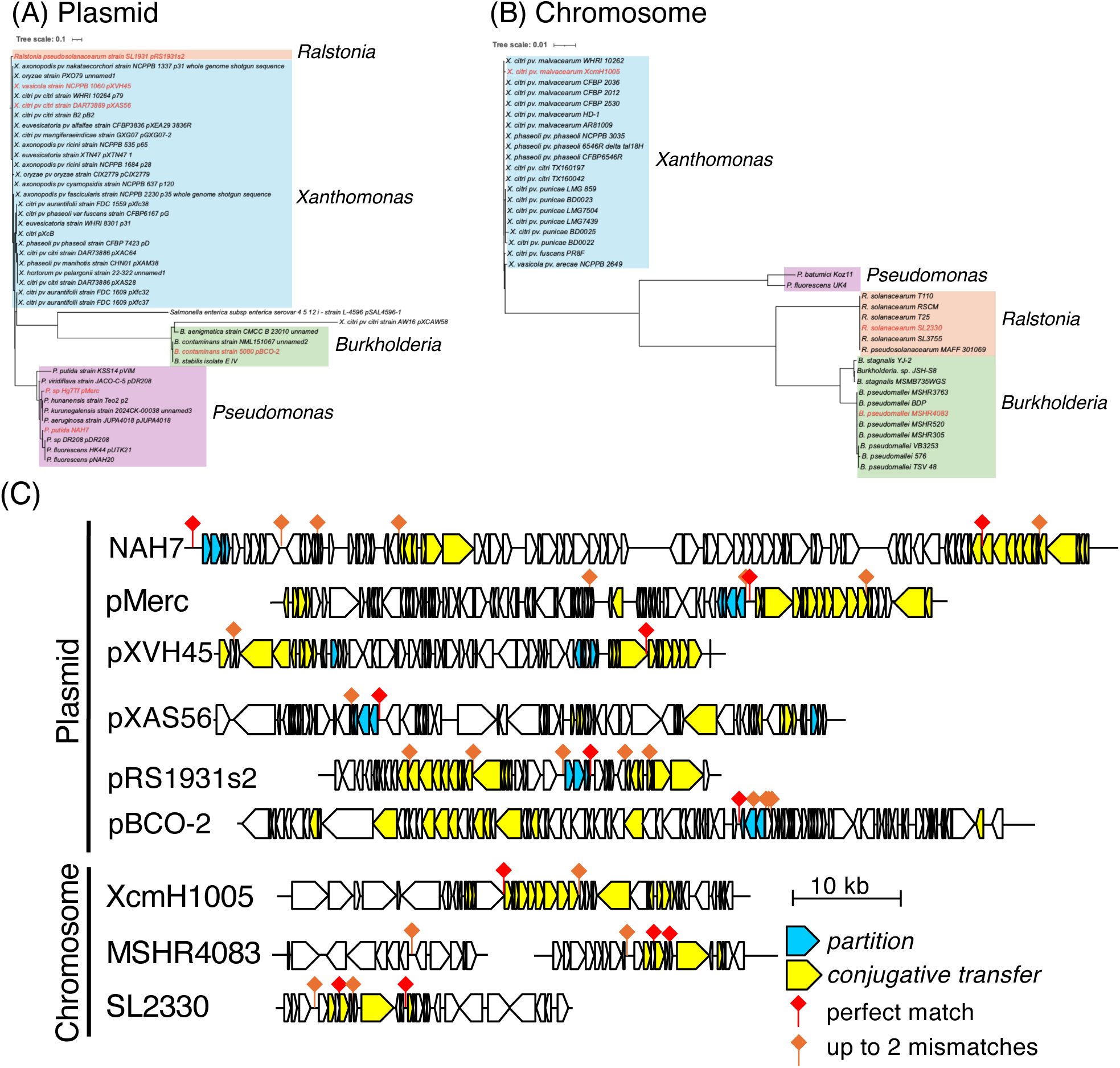
Phylogenetic analysis and distribution of *parS_NAH_* sequences in plasmid and chromosomal sequence. **(A, B)** Phylogenetic trees of sequences containing *parS_NAH_* sites. The sequences containing perfectly matching *parS1_NAH_* were identified through BLAST searches. The plasmid-associated tree (A) was constructed using amino acid sequences of relaxase, while the chromosomal tree (B) used 16S rRNA gene sequences. Both trees were constructed using MAFFT for sequence alignment and visualized with iTOL. The sequences are categorized into the genera *Pseudomonas*, *Burkholderia*, *Xanthomonas*, and *Ralstonia*. **(C)** Schematic representation of the genomic context surrounding *parS1_NAH_* sequences and those with up to two nucleotide mismatches in selected plasmids and chromosomal regions. The *parS1_NAH_* sites are indicated by red diamonds (perfect matches) and orange diamonds (up to 2 mismatches). Genes involved in partitioning (cyan) and conjugative transfer (yellow) are highlighted. The schematic represents the approximate locations of these elements within the 10 kb genomic regions surrounding the *parS1_NAH_* sites. For plasmids, the entire sequence length is represented, while for chromosomes, only the surrounding regions containing the repeat sequences are shown. Specifically, for chromosomal sequences: XcmH1005 (GenBank: CP013004) represents from positions 187,208 to 227,688. MSHR4083 (GenBank: CP017050) represents from positions 1,091,874 to 1,109,185 and 1,954,109 to 1,971,627.SL2330 (GenBank: CP022794) represents from positions 594,289 to 620,391.

In addition, we selected representative plasmid and chromosomal sequences to assess the presence of *parS_NAH_* variants. This analysis identified not only perfectly conserved *parS1_NAH_* but also those permitting up to two mismatches. The schematic representation (Fig. 4C) illustrates the distribution of *parS_NAH_* sites, highlighting their positions relative to conserved partitioning and conjugative transfer genes. Importantly, their proximity to partitioning and conjugative transfer genes suggests a potential functional link between these elements and the regulation of plasmid stability and horizontal gene transfer.

### Binding profiles of ParR, TraA, and TraD to the *oriT* region

The role of ParR in activating the transcription of conjugation-related genes has not been supported by experimental evidence. Since ParR contains a KORA domain predicted to bind DNA, its DNA-binding activity likely contributes to the observed increase in conjugation frequency. In the process of conjugative transfer, *oriT* is a particularly important DNA region, serving as the initiation site for DNA transfer. The binding of proteins to this region is believed to play a critical role in facilitating conjugation. Based on this, we investigated whether ParR specifically binds to the *oriT* region to explore its potential role in conjugative transfer (Fig. 5). The EMSA demonstrated that ParR binds to the *oriT* DNA, forming multiple distinct complexes (Fig. 5A). The disappearance of the free probe and the appearance of shifted bands confirm the interaction between ParR and the DNA. Notably, ParR produced a greater number of shifted bands, suggesting the formation of diverse binding complexes.

**Fig. 5.**
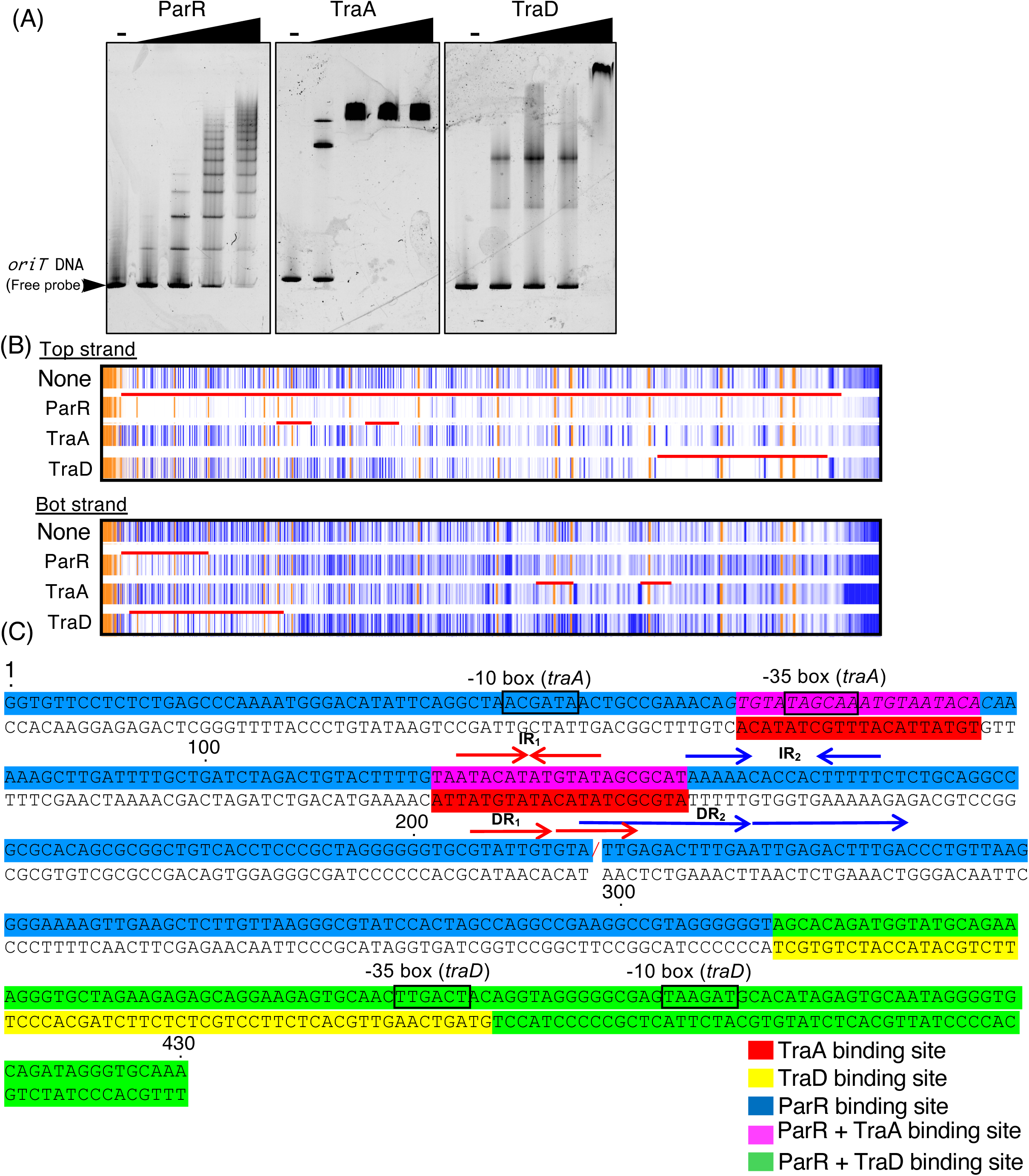
Protein-DNA interactions of ParR, TraA, TraD with *oriT* DNA. **(A)** EMSA demonstrating the binding of ParR-Strep, TraA-His, and TraD-His proteins to *oriT* DNA fragments. The *oriT* DNA fragment (9 nM) was incubated with ParR-Strep, TraA-His, and TraD-His at final concentrations of 0.1, 0.2, 0.5, and 0.9 µM. The binding reactions were performed at 30°C for 5 minutes and analyzed using 15% polyacrylamide gel electrophoresis with Tris-glycine buffer. The gels were stained with SYBR Gold and visualized using the FLAS-T1500 system. The "*oriT* DNA" represents the free DNA probe. **(B)** DNase I foot-printing assay results showing the protection pattern on *oriT* DNA by ParR-Strep, TraA-His, and TraD-His proteins. The assay was conducted using FAM-labeled *oriT* DNA fragments to identify the binding regions on both the top and bottom strands. The orange regions represent internal standard markers (GeneScan 500 LIZ size standard), while the blue lines indicate the FAM-labeled *oriT* DNA fragments obtained after DNase I treatment. DNase I-protected regions indicate protein-DNA interaction sites, with red line highlighting specific binding regions compared to the ‘None’ control. The data were analyzed using TraceViewer, with corresponding peak data available in Fig. S3 and S4 (C) Sequence analysis of the *oriT* region, highlighting the binding sites for ParR, TraA, and TraD proteins. This panel was created based on foot-printing analysis (panel B) and G+A ladder data from Fig. S3 and S4. Specific binding sites are color-coded as follows: ParR binding sites are indicated in blue, TraA binding sites in red, and TraD binding sites in orange. Overlapping binding sites between ParR and TraA are shown in purple, while those shared by ParR and TraD are marked in green. "/" indicates the *nic* site. The −10 and −35 boxes are indicated, representing promoter regions relevant to the transcriptional regulation of conjugative transfer genes. Important IR and DR sequences identified in previous studies as significant for their functions are also annotated.

Homologs of TraA and TraD—specifically TrwA and StbA in the well-studied R388 plasmid—are known to bind the *oriT* region (4, 9). To determine whether TraA and TraD similarly interact with the *oriT* DNA in the NAH7 system, we performed EMSAs (Fig. 5A). Both TraA and TraD bound to the *oriT* region, as indicated by the disappearance of the free probe and the appearance of shifted bands. TraA and TraD displayed a clear binding pattern characterized by discrete, well-defined shifted bands at all tested concentrations.

We further examined ParR’s DNA-binding profile through DNase I foot-printing with a FAM-labeled *oriT* DNA fragment. As shown in Fig. 5B, ParR extensively protected the top strand of the *oriT* DNA, including the *nic* site, a key element in conjugative transfer (Fig. 5C). This widespread protection suggests that ParR engages multiple sites along the DNA, possibly through cooperative binding. On the bottom strand, ParR’s protection was more localized, indicating asymmetric binding or varying accessibility between strands. TraA protected the −35 box of the *traABC* gene cluster promoter region and the IR1 sequence (Fig. 5B), which are essential for conjugative transfer (17). TraD, in contrast, protected a broader region encompassing the promoter of the *traDEF* gene cluster, including both the −35 and −10 boxes.

## Discussion

As part of elucidating the molecular mechanism of conjugative transfer activation in NAH7, this study investigated the role of the *par* gene cluster, which contributes to plasmid stability (Fig. 1D), in enhancing conjugative transfer (Fig. 2). Our findings demonstrated that ParB acts as a transcriptional activator for genes encoding components of the T4SS, while ParR increases the frequency of conjugative transfer, likely by binding to the *oriT* region, without directly activating the transcription of conjugation-related genes (Fig. 6). Based on the mechanism of action of ParB and ParR in enhancing the conjugative transfer frequency of pKKO3-2, it is that these proteins also enhance the conjugative transfer frequency of the native NAH7 plasmid. In NAH7-harboring strains, a slight increase in conjugative transfer frequency was observed upon the expression of either *parB* or *parR* (Fig. S5). However, the presence of *par* genes on NAH7 may mask the frequency increase, making it challenging to detect further enhancements. In contrast, ParA seems to enhance conjugative transfer by increasing the copy number of the original vector, pNIT6012, derived from pKKO3-2 (Fig. 3G and H). This increase in copy number likely leads to elevated transcription levels of conjugation-related genes, resulting in a higher frequency of conjugative transfer. No evidence was found to support a direct role for ParA in enhancing the conjugative transfer frequency of the native NAH7 plasmid.

**Fig. 6.**
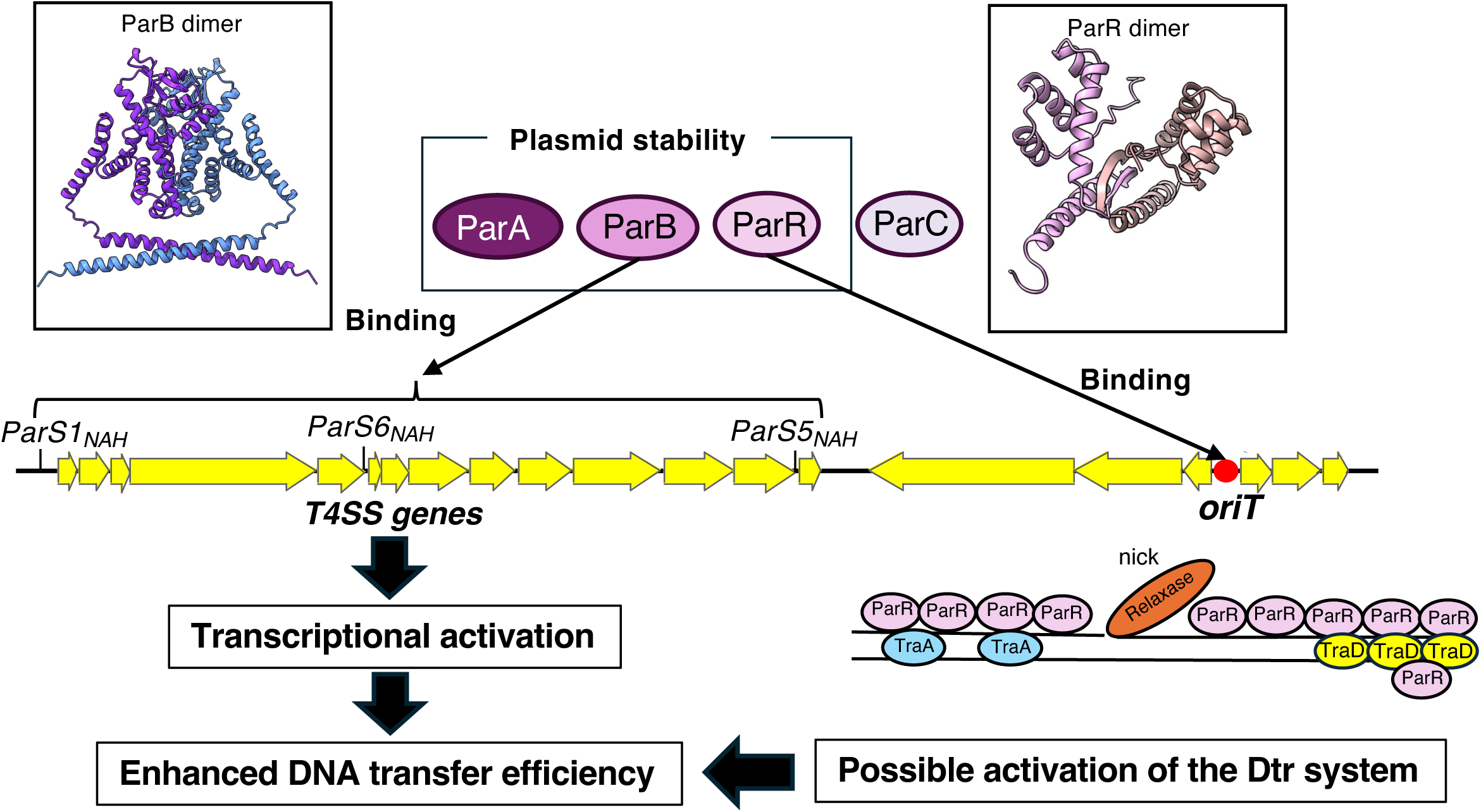
Proposed model of Par proteins in conjugative transfer enhancement in the NAH7 system. This model illustrates the functional roles of ParA, ParB, ParR and ParC proteins in plasmid stability and conjugative transfer in the NAH7 system. ParA, ParB, and ParR contribute to maintaining plasmid stability within host cells Structures of ParB and ParR are depicted as dimers, highlighting their roles in binding DNA. ParB binds specifically to *parS_NAH_* sites (*parS1_NAH_*, *parS5_NAH_*, and *parS6_NAH_*), which are located near the T4SS genes (yellow arrows). The binding of ParB to *parS* sites results in the transcriptional activation of T4SS-related genes, promoting the assembly of the T4SS and thus enhancing DNA transfer efficiency. At the *oriT* region, ParR binds specifically, potentially aiding in the recruitment of conjugative machinery and interacting with relaxase and T4SS components, such as TraA and TraD. This interaction may help facilitate the initiation of DNA transfer, promoting the Dtr system during conjugation.

### Biological significance of *par* genes in conjugative transfer enhancement

Since plasmid partitioning is synchronized with host cell division, under nutrient-rich and favorable conditions that support host cell division, probably NAH7 not only efficiently distributes to daughter cells but also activates conjugative transfer, enabling its spread to other bacterial cells. This dual functions may represent an adaptive strategy for the plasmid, as its original host, *Pseudomonas putida* G7, typically inhabits nutrient-poor soil environments (27). The ability to transfer to other cells under nutrient-rich conditions could provide a significant survival advantage for the plasmid.

In Gram-negative bacteria, the promoters of conjugation-related genes are often regulated by negative feedback mechanisms involving transcriptional repressors, which is well-understood in the context of zygotic induction (28) (29). For the NAH7 plasmid, transfer begins at the *oriT* site and proceeds counterclockwise on the plasmid map (Fig. 1A) (17). The *par* gene cluster, located on the T-strand, is transferred before the T4SS-related genes. This ensures the immediate expression of ParB in the recipient cell, which in turn activates the transcription of T4SS-related genes. The early expression of ParB enables the recipient cell to quickly transition into a donor, facilitating further plasmid dissemination. Foot-printing analysis showed that ParR specifically binds to the T-strand (the top strand in Fig. 5B), suggesting that ParR may also function to protect the T-strand within the recipient cell from enzymes that cleave DNA. Moreover, other plasmid systems have demonstrated that certain Par proteins can be transferred to the recipient cell via T4SS (14, 30–32). ParB and ParR might also be transferred and contribute to enhancing conjugative transfer activity within the recipient cell.

### Mechanisms of enhanced conjugative transfer by ParR and ParB in NAH7: comparison with KorA and KorB in RP4

In the IncP-1α plasmid RP4, the CBP KorB acts as a transcriptional repressor, specifically targeting the *trbB* promoter, and this repressive action is further enhanced by cooperative interactions with another repressor TrbA (33). In contrast, our study demonstrated that ParB binds to the *parS_NAH_* sites and promotes the transcription of conjugation-related genes (*mpfA*, *mpfE*, and *mpfK*) located downstream of this site. The similarity between ParB and KorB is low, particularly in the DNA-binding domain (Fig. S6B). Although both the *parS* site in NAH7 and the OB sites in RP4 are IR sequences (Fig. S6A), their lengths and sequences differ significantly, reflecting their distinct functional roles—transcriptional activation by ParB versus repression by KorB. Additionally, KorB is known to work cooperatively with other DNA-binding proteins such as KorA to achieve effective transcriptional repression (34). While it remains unclear whether ParB also collaborates with other regulatory factors, our findings indicate a potential cooperative interaction between ParB and ParR. Co-expression of ParB and ParR led to a higher conjugative transfer frequency of pKKO3-2 compared to the expression of either protein alone.

ParR, a homolog of KorA, is known to function as a transcriptional repressor. However, no evidence supports ParR regulates to conjugation-related genes. Instead, EMSA and DNase I foot-printing assays revealed that ParR binds to multiple sites on the *oriT* DNA with varying affinities or forms oligomers with different stoichiometries. The ParR binding to *oriT* DNA may play a role in facilitating its function in conjugative transfer. It is also possible that ParR, in interactions with the relaxase proteins or TraA, contributes to the assembly of the relaxosome, thereby enhancing the efficiency of conjugative transfer. ParR likely has functions beyond binding to *oriT*, as the disruption of the *parR* gene led to reduce plasmid stability even in the absence of *oriT* (e.g., in pK7OR). This suggests that ParR contributes to plasmid stability through *oriT*-independent mechanisms.

### Unexpected increase in plasmid copy number induced by ParA

ParA produced an unexpected result by increasing the copy numbers of the vectors pNIT6012 and pKKO3-2. The vector pNIT6012 contains both the p15a origin of replication, which operates with the replication machinery in *E. coli*, and the broad-host-range replication system pVS1. It is believed that ParA acts on one of these systems to promote an increase in copy number. The ability to control plasmid copy number is considered valuable as genetic tools (35, 36), especially since plasmids transferred through conjugation play a crucial role in genetic engineering and biotechnology.

### Comparison with other conjugative transfer enhancement mechanisms

Mechanisms involving the enhancement of conjugative transfer by partitioning proteins have also been reported in other plasmids, such as plasmid R1 and the Ti plasmid (14, 37). In plasmid R1, the Type II partition system ATPase ParM_R1_ and the centromere-binding protein ParR_R1_ interact directly with components of the conjugative machinery, thereby enhancing DNA transfer efficiency. Specifically, ParM_R1_ and ParR_R1_ physically and functionally interact with the relaxase, stimulating its relaxase activity, which is essential for initiating DNA transfer. Additionally, ParM_R1_ interacts with T4SS ATPases such as TraD _R1_ (VirD4) and TraC _R1_ (VirB4), indicating a coordinated role in facilitating the assembly and function of the T4SS. In contrast, in the NAH7 system, we found that conjugative transfer frequency increases through the binding of ParB or ParR to DNA, while the involvement of direct protein-protein interactions remains unclear. Using the AlphaFold3 server, we predicted the interactions between six protein pairs: ParA, ParB, and ParR with TraC (relaxase), TraB (VirD4), and MpfC (VirB4). All ipTM (interface predicted TM-score), which is used to evaluate the confidence of predicted relative positions of subunits forming a protein-protein complex, values obtained were relatively low, with the highest interaction scores being observed for ParA - MpfC, ParA - TraC, and ParR - TraC, each around 0.35. Given that the highest ipTM value in our analysis was 0.35, these predictions indicate a low likelihood of stable interactions between the tested protein pairs. Since NAH7 and R1 possess different types of partitioning systems, it is less likely that NAH7 utilizes a mechanism similar to R1 involving direct protein-protein interactions for enhancing conjugative transfer frequency. Instead, the enhancement in NAH7 may rely on different molecular strategies, possibly involving DNA-binding rather than direct interactions with the conjugative machinery.

## MATERIALS AND METHODS

### Strains and growth conditions

Bacterial strains listed in Table 1 were grown in Lysogeny Broth (LB). *P. putida* KT2440 strains were grown in one - third LB media for plasmid stability assay 30°C. Strains and plasmid were maintained with appropriate antibiotic selection: ampicillin (100 μg/ml), tetracycline (20 μg/ml), kanamycin (50 μg/ml), and rifampicin (100 μg/ml).

**Table 1.**
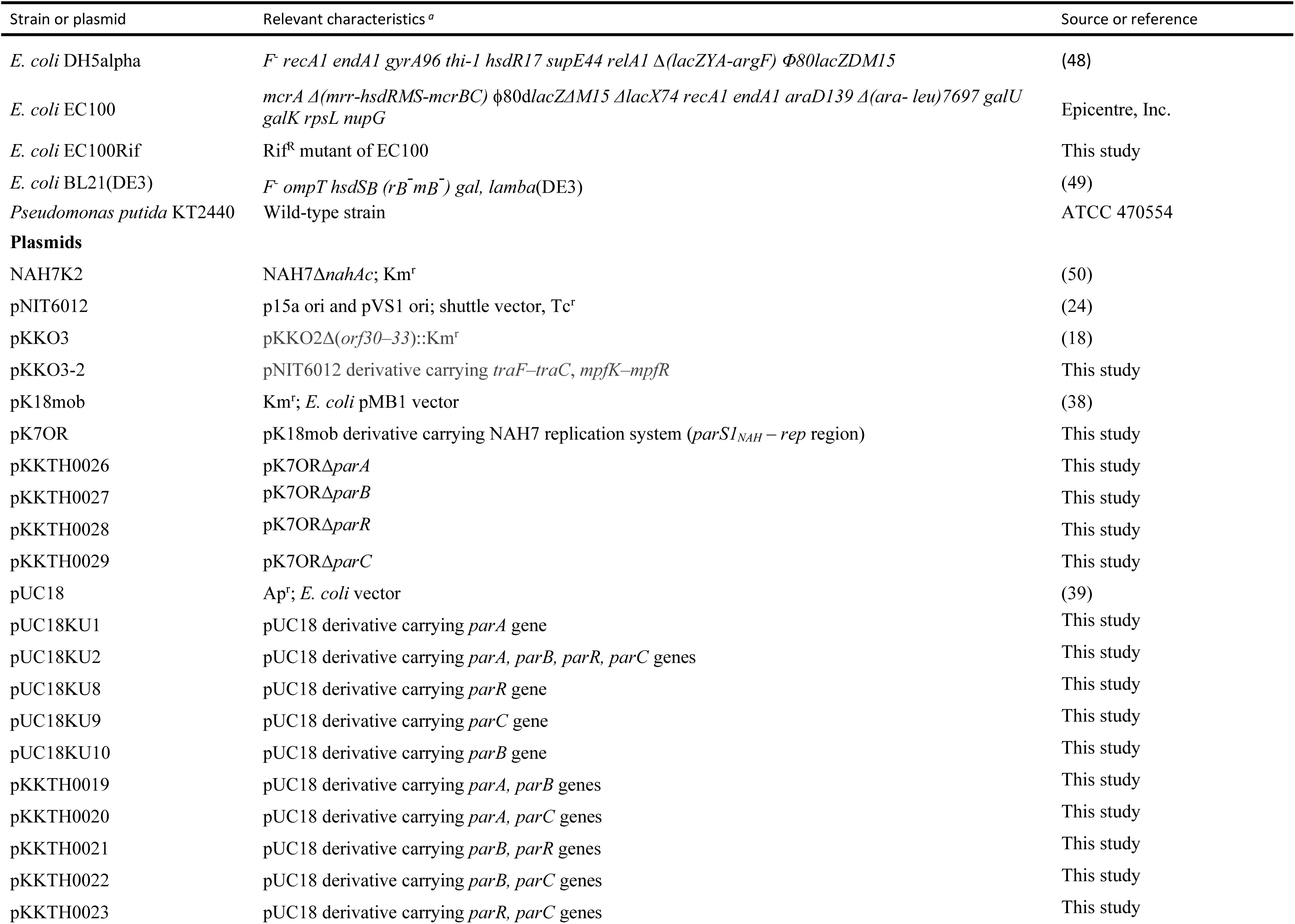
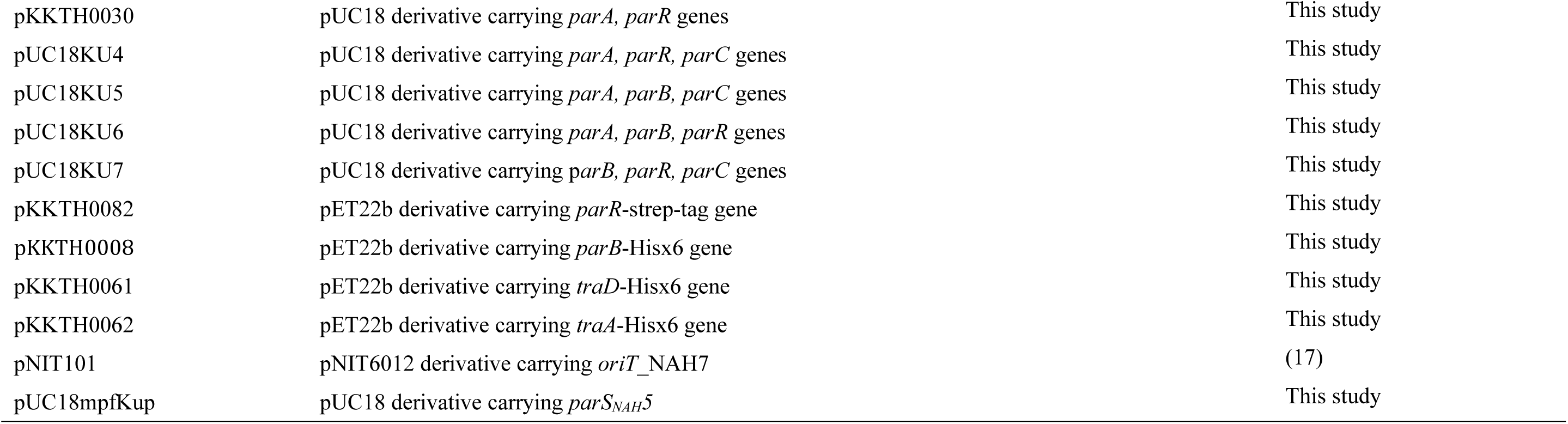
Bacterial strains and plasmids used in this study.

### Plasmid constructions

All plasmids and oligonucleotide primers used in these studies are listed in Tables 1 and S1. pKKO3-2 is a plasmid derived from the self-transmissible plasmid pKKO3 (18) with the Km resistance gene removed. The Km resistance gene on pKKO3 is flanked by KpnI sites, allowing it to be excised by KpnI digestion, followed by self-ligation. The removal of the Km resistance gene was confirmed by PCR and by the loss of Km resistance.

pK7OR is a mini-NAH7 plasmid containing the region from *parS1_NAH_* to the *rep* gene. A DNA fragment containing this region was amplified from NAH7, digested with XhoI and SpeI, and inserted into the SalI - XbaI site of pK18mob (38). Plasmids pKKTH0026 to pKKTH0029 are derivatives of pK7OR, each lacking one of the *par* genes. To construct these plasmids, DNA fragments with specific *par* gene deletions were amplified from pK7OR and self-ligated using the Gibson Assembly kit (New England Biolabs, Beverly, MA).

pUC18KU1, pUC18KU8, pUC18KU9 and pUC18KU10, each expressing a single *par* gene, were constructed by amplifying individual *par* gene regions from NAH7 and cloning them into the KpnI site of pUC18 (39) using the Gibson Assembly kit. Plasmid pUC18KU2, which expresses all four *par* genes, was similarly constructed by amplifying the entire *par* gene region from NAH7 and cloning it into the KpnI site of pUC18 using the Gibson Assembly kit.

Plasmids expressing combinations of two *par* genes were constructed by amplifying DNA fragments from specific templates and cloning each fragment into the KpnI site of pUC18 using the Gibson Assembly Kit. For pKKTH0019, pKKTH0021, pKKTH0022, pKKTH0023, and pKKTH0030, the respective templates used were pUC18KU2, pUC18KU2, pUC18KU5, pUC18KU2, and pUC18KU4. Plasmid pKKTH0020 was constructed by amplifying a fragment from pUC18KU4, phosphorylating it, and performing self-ligation.

Plasmids expressing combinations of three *par* genes were constructed by amplifying specific gene regions from NAH7 and cloning each combination into the KpnI site of pUC18 using the Gibson Assembly Kit. For pUC18KU6 and pUC18KU7, the *parA-parR* and *parB-parC* regions were amplified with primer sets KUDO011-KUDO29 and KUDO012-KUDO038, respectively. For pUC18KU4 and pUC18KU5, the *parA* and *parRC* regions, and the *parAB* and *parC* regions, were amplified using primer sets KUDO011-KUDO026, KUDO012-KUDO025, KUDO011-KUDO028, and KUDO012-KUDO027, respectively. Each construct was designed to be IPTG-inducible.

Plasmids pKKTH0082, pKKTH0008, pKKTH0061, and pKKTH0062 were constructed to express affinity-tagged proteins. DNA fragments of each gene region were amplified and cloned into the NdeI and XhoI sites of pET22b using the Gibson Assembly Kit. For pKKTH0082, which expresses ParR-strep, the oligonucleotide primer (oligoNucKK0319) included a sequence for the Strep-tag.

The plasmid pUC18mpfKup, containing the upstream *parS5_NAH_* region of *mpfK*, was constructed by annealing two oligonucleotides, Bam_up_orf34_kpn and Bam_up_orf34_kpn_COM, to create a double-stranded DNA fragment. This fragment was then digested with BamHI and KpnI and ligated into the corresponding sites of pUC18.

### Plasmid stability assay

pK7OR and its derivative plasmids, which contain the Km resistance gene, were tested for stability using *P. putida* KT2440 as the host strain. The strains were first grown overnight in one - third LB medium supplemented with kanamycin. Subsequently, cultures were diluted 0.1% into fresh one - third LB medium without antibiotics and incubated for 24 hours. After incubation, the cultures were plated onto one - third LB agar plates without antibiotics. On the following day, 100 colonies were randomly picked and streaked onto one - third LB agar plates with kanamycin. The proportion of Km-resistant colonies was then calculated. The assay was performed in three independent biological replicates. The average values were plotted with individual data points and standard deviations represented as error bars.

### Conjugation assay

Donor and recipient cells were grown overnight at 30°C in the presence of appropriate antibiotics. Each culture (1 mL) was centrifuged to harvest the cells, and the resulting pellets were resuspended in 50 µL of fresh medium. The suspensions were combined to a final volume of 100 µL and spotted onto LB agar plates supplemented with 0.5 mM IPTG for induction, then incubated for 6 hours at 30°C. Following incubation, cells were resuspended in 1 mL of LB, serially diluted in LB, and plated on LB agar containing antibiotics to select for transconjugants (Tc), recipients, and donors (D). Plates were incubated overnight at 30°C. DNA transfer frequency is reported as the number of transconjugants per donor (Tc/D). All mating experiments were performed at least three times in triplicate, and representative data is shown with individual data points, average transfer frequencies as bars, and standard deviations as error bars.

### Quantification of the expression of conjugative transfer genes in pKKO3-2

RNA extraction samples were prepared as follows: pKKO3-2 strains expressing *par* genes from pUC18 were each grown overnight, and 1 mL of each culture was centrifuged to harvest the cells. The cell pellets were resuspended in 50 µL of fresh medium. This culture was spotted onto LB agar plates containing 0.5 mM IPTG and incubated at 30°C for 3 hours. After incubation, the cells were resuspended in LB medium, and RNA was extracted from these samples using RNeasy (QIAGEN, Hilden, Germany) following the manufacturer’s protocol.

After extraction, each RNA sample was treated with DNase I (TAKARA, Shiga, Japan) at 37°C for 2 hours to remove residual DNA. Reverse transcription was performed with ReverTra Ace qPCR RT Master Mix (Toyobo, Osaka, Japan) and random 9-mer primers. qRT-PCR was conducted using Luna Universal qPCR Master Mix (New England Biolabs, Beverly, MA) on a CFX Connect Real-Time System (Bio-Rad Laboratories, Hercules, CA). The PCR protocol included an initial denaturation at 95°C for 1 minutes, followed by 40 cycles of 10 seconds at 95°C, 10 seconds at 60.0°C, and 10 seconds at 68°C.The primer sets oligoNucKK0074-oligoNucKK0075, oligoNucKK0076-oligoNucKK0077, oligoNucKK0078-oligoNucKK0079, oligoNucKK0080-oligoNucKK0081, oligoNucKK0082-oligoNucKK0083, oligoNucKK0106-oligoNucKK0107, and oligoNucKK0084-oligoNucKK0085 were used to quantify *mpfA*, *mpfE*, *mpfK*, *traD*, *traB*, *tetA* and *gyrA*, respectively. Expression levels of each target gene were normalized to the expression level of the *gyrA* gene on the *E. coli* chromosome. Individual data points, average expression levels are represented as bars, and standard deviations are shown as error bars.

### Electrophoretic mobility shift assay (EMSA)

ParB-His, TraA-His, TraD-His, and ParR-Strep were expressed in BL21(DE3) cells containing the expression plasmids pKKTH0008, pKKTH0062, pKKTH0061, and pKKTH0082, respectively. Cells were cultured at 30°C, and protein expression was induced at an OD660 of approximately 0.5 by adding 0.5 mM IPTG. After 6 hours induction, cells were harvested and resuspended in RIPA buffer (50 mM Tris-HCl (pH 8.0), 150 mM NaCl, 1% (v/v) Triton X-100, 1% (w/v) sodium deoxycholate, and 0.1% (w/v) SDS). Following sonication, the supernatant was used for protein purification. His-tagged proteins were purified using Ni-NTA agarose (Qiagen, Hilden, Germany), and the Strep-tagged protein was purified using the Strep-Tactin XT 4Flow Starter Kit (IBA, Göttingen, Germany). The purity of each protein was confirmed by SDS-PAGE and CBB staining (Fig. S7).

The purified proteins, TraA-His, TraD-His, and ParR-Strep, were incubated with the *oriT* DNA fragment (9 nM) in a binding buffer containing 100 mM NaCl, 80 mM Tris-HCl (pH 7.5), 0.1 mM EDTA, and 5 mM MgCl₂ at 30°C for 5 minutes. Final protein concentrations of 0.1, 0.2, 0.5, and 0.9 µM were prepared for these experiments.

For ParB-His, a different binding buffer was used, composed of 50 mM Tris-HCl (pH 7.5), 10 mM MgCl₂, 1 mM ATP, and 10 mM DTT. The ParB-His protein was incubated with DNA corresponding to the upstream *parS5_NAH_* region of *mpfK* at a final DNA concentration of 23 nM at 30°C for 15 minutes. The final concentrations of ParB-His were 0.15, 0.3, 0.45, and 0.9 µM. Samples were electrophoresed on a 15% polyacrylamide gel at 200 V for 90 minutes using Tris-glycine buffer, stained with SYBR Gold (Invitrogen, Carlsbad, CA, USA), and detected using the FLAS-T1500 system (Fujifilm, Tokyo, Japan).

### DNase I Foot-printing Assay

Two types of FAM-labeled *oriT* DNA fragments were amplified using pNIT101 as the template with primer sets pNIT5041_FAM-pNIT5548 and pNIT5041-pNIT5548_FAM, ensuring that each fragment was labeled on a single strand. The purified proteins TraA-His and TraD-His were incubated with the *oriT* DNA fragment (12 nM) at a final concentration of 1 µM each, and ParR-Strep was incubated at a final concentration of 8 µM. The binding reactions were performed in a buffer containing 100 mM NaCl, 80 mM Tris-HCl (pH 7.5), 0.1 mM EDTA, 5 mM MgCl₂, and 50 µg/mL salmon sperm DNA at 30°C for 15 minutes. Following the incubation, the reaction mixtures were treated with 1/10 unit of DNase I at 12°C for 1 minutes. The reactions were stopped by heating at 98°C for 10 minutes to inactivate DNase I. The DNase I-treated fragments were purified using the BigDye XTerminator purification kit (Thermo Fisher Scientific).

The purified products were mixed with the GeneScan 500 LIZ size standard (Life Technologies) and separated using an ABI Prism 3130xl Genetic Analyzer. To prepare a G+A sequencing ladder sample, the method of Eckert (40) was used, treating the FAM-labeled dsDNA fragments with formic acid and piperidine. The resulting data were analyzed using the TraceViewer software (41). The raw data supporting these findings are presented in Fig. S3 and Fig. S4.

### Bioinformatics Analysis

To estimate the functions of the Par proteins from the NAH7 plasmid (Fig. S1), the amino acid sequences were used as queries for InterPro analysis to identify conserved domains and predict functional annotations. To obtain predicted structures of these Par proteins, we utilized the AlphaFold Database (42) and AlphaFold Server (43) for structure prediction.

To assess the conservation of ParABRC homologs of NAH7 within IncP-9 plasmids (16), a BLASTP analysis was conducted using the protein sequences of IncP-9 plasmids as a database. Each of NAH7’s ParA, ParB, ParR, and ParC proteins was used as a query. An E-value threshold of 10^-8^ was applied to determine significance. Hits meeting this criterion were marked as "present," whereas sequences without significant hits were labeled as "absent." The results are summarized in Table S2.

To identify plasmid and chromosomal sequences that perfectly match the NAH7 *parS1* site (18 bp sequence: 5′-TTTCTCGCATGCGAGAAA-3′), we used the NCBI Dataset tool to filter for "complete plasmid" and "complete genomes" datasets. The filtered sequences were then subjected to BLASTN searches to detect the presence of the *parS1_NAH_* site across different bacterial genomes. To classify the identified sequences phylogenetically, we utilized the amino acid sequences of plasmid-encoded relaxase genes and the DNA sequences of 16S rRNA genes from chromosomes. Detection of relaxase genes was performed by conducting a BLASTP search against the experimentally verified relaxase sequences obtained from oriTDB (44) with an e-value cutoff of 1e-5. If multiple plasmids carried identical relaxase genes, we selected a single representative sequence from among them for all subsequent analyses. The 16S rRNA gene sequences were extracted from GenBank by selecting annotated 16S rRNA entries. The detected sequences were aligned using MAFFT (45) with NJ method, and a phylogenetic tree was constructed to visualize the relationships among the sequences. The resulting tree was modified for better visualization using iTOL (46).

From representative sequences, EMBOSS: fuzznuc was used to find *parS1_NAH_* site that perfectly matched or had up to two mismatches. The approximate positions of these sites were plotted on gene maps in PowerPoint for visual representation.

### Statistical analysis

All statistical analyses were performed using GraphPad Prism 9. Ordinary one-way ANOVA with Dunnett’s multiple comparisons test or unpaired two-tailed t-tests were used as specified in the figure legends. P-values and statistical significance are indicated in the figures.

## ACKNOWLEDGMENTS

This work was supported by Grant-in-Aid for Early-Career Scientists (Grant ID: 19K15725) and Institute for Fermentation, Osaka (IFO) (Grant ID: Y-2024-1-007) Ko.Ki. performed most of the experiments and drafted initial manuscript. Ko.Ku., Re.Ku., We.De. performed several conjugation assays. Ko.Ki. and Ma.Ts. supervised the whole project. All authors, Ko.Ki., Ko.Ku., Re.Ku., We.De., Na.Ki., Le.St., Yo.Oh, Y.N., and M.T., contributed to discussions, editing, and revision of the manuscript.

## Supporting Information

**Figure S1.**
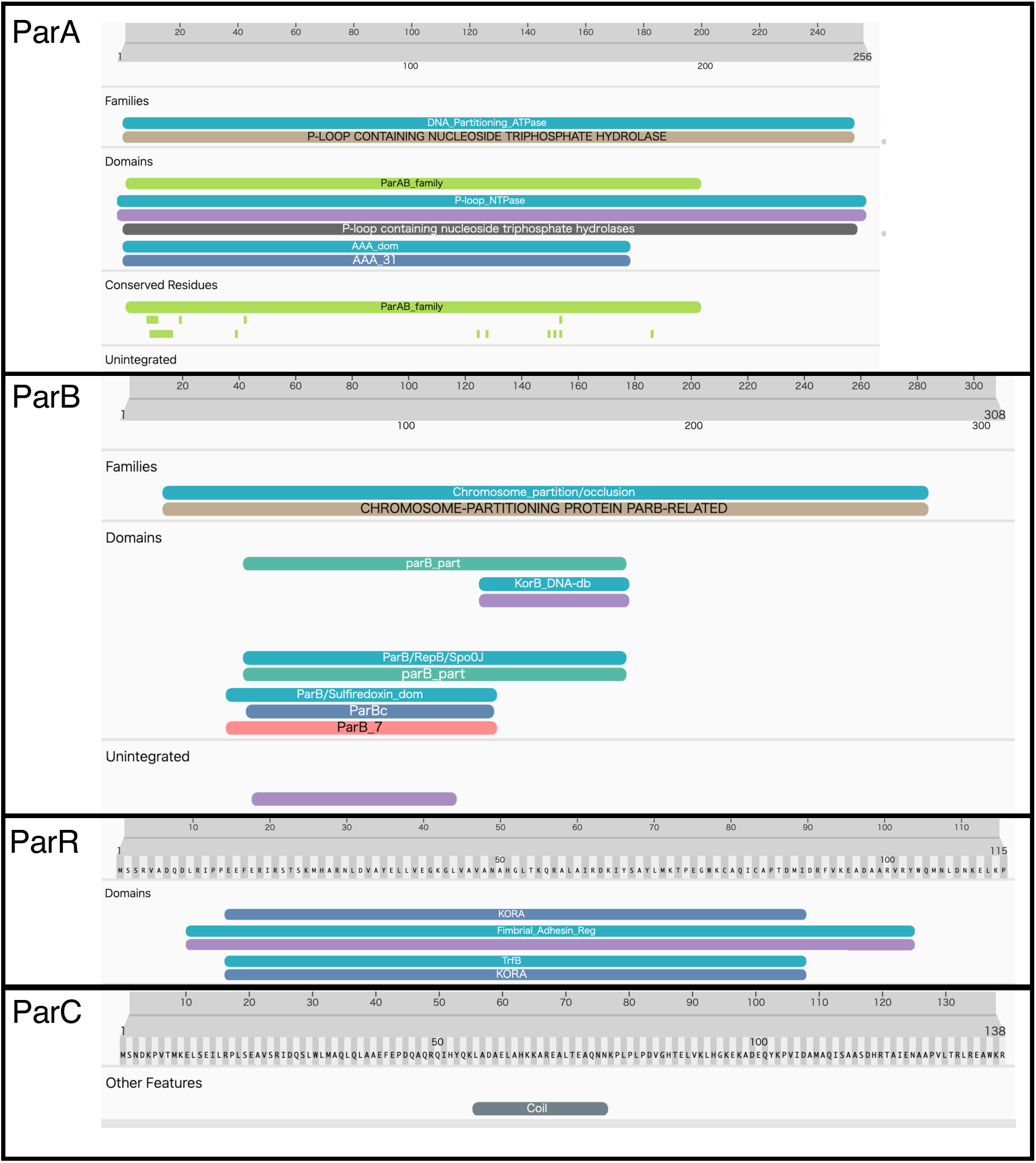
Domain architecture of NAH7 Par proteins as predicted by UniProt Analysis. This figure illustrates the predicted domain structures of ParA, ParB, ParR, and ParC. Domains were identified using UniProt and are color-coded to indicate different functional components.

**Figure S2.**
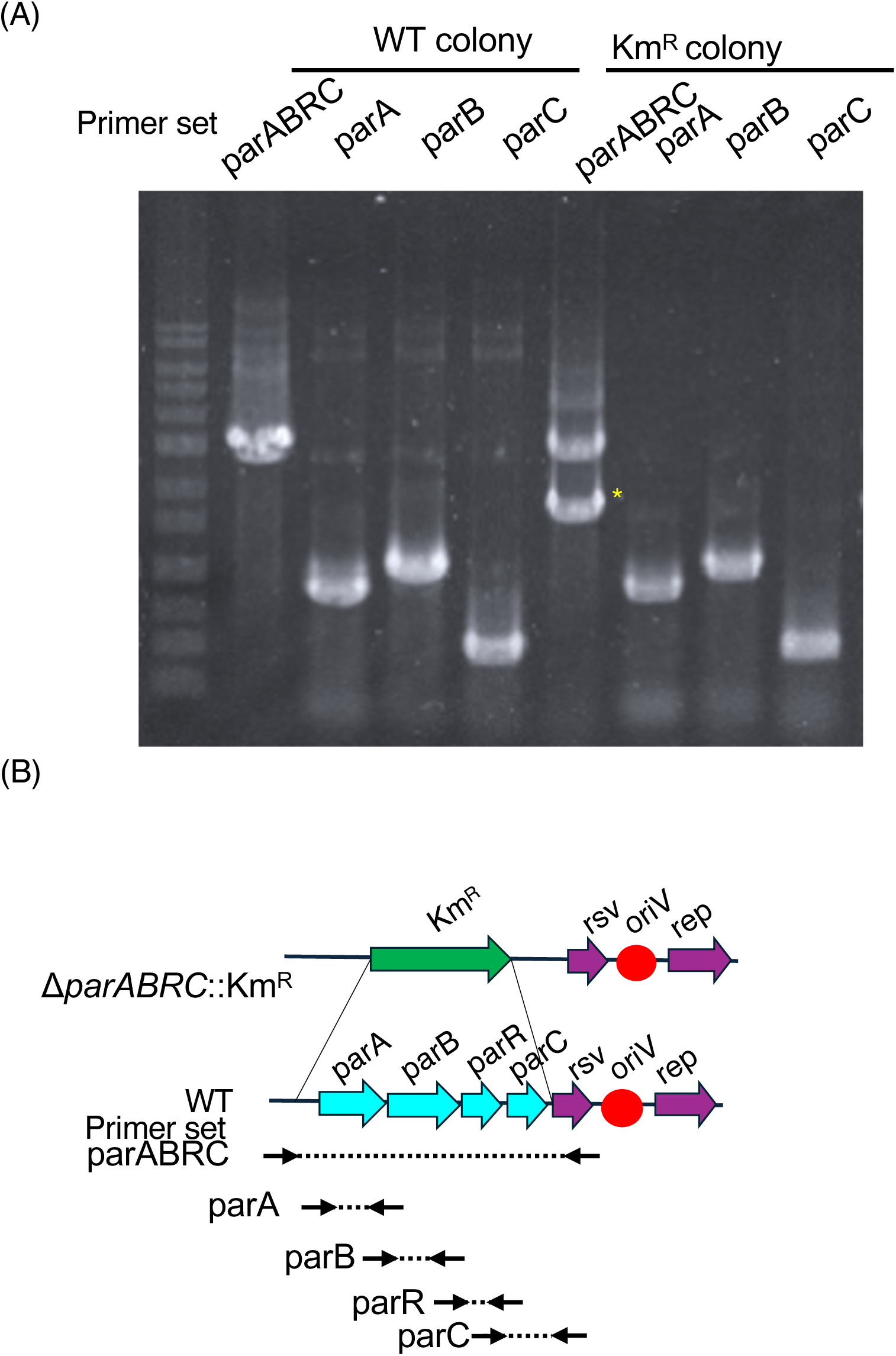
Inability to isolate a pure NAH7Δ*parABRC*::*Km^R^* mutant. **(A)** Colony PCR analysis comparing wild-type (WT) NAH7 and colonies presumed to carry a Δ*parABRC*::*Km^R^* plasmid. The primer sets (labeled above each lane) target the *parABRC*, *parA*, *parB*, and *parC* regions. The asterisk (*) marks the DNA fragment replaced by the Km cassette. Despite obtaining Km-resistant colonies, PCR consistently shows both the WT par genes and the Km^R^ fragment, indicating a mixed population of WT and Δ*parABRC*::*Km^R^* plasmids in the same colony. (B) Schematic diagram of the *parABRC* region in NAH7. The cyan arrows represent *parA*, *parB*, *parR*, *parC*, and the green arrow depicts the *Km^R^* cassette replacing the *parABRC* locus. The dashed arrows indicate the approximate positions of the PCR primer sets. These results suggest that the *par* genes are required for stable maintenance of NAH7, preventing isolation of a pure Δ*parABRC*::*Km ^R^* mutant plasmid.

**Figure S3.**
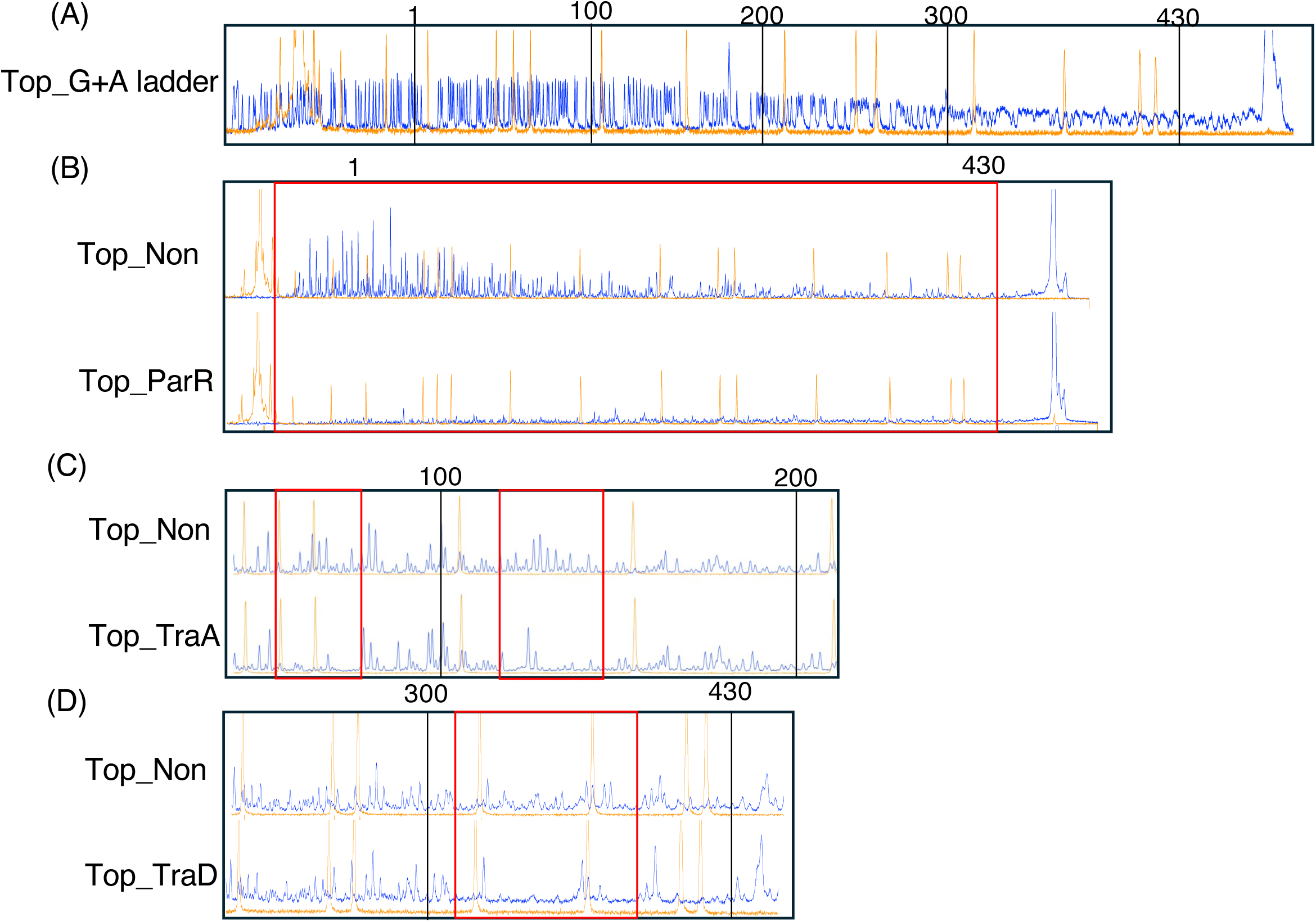
DNase I foot-printing analysis of the *oriT* region on the top strand. This figure illustrates the results of DNase I foot-printing assays conducted to identify protein binding regions along the top strand of the *oriT* DNA. The blue peaks represent DNA cleaved by DNase I, and the orange peaks represent internal standards for alignment. (A) The G+A ladder serves as a reference to locate specific nucleotide positions on the DNA strand, corresponding to the *oriT* region depicted in Fig. 5(C), enabling precise mapping of protein-DNA interactions. (B) Foot-printing results with PraR-strep protein. The non-protein control ("Top Non") shows the baseline cleavage pattern, and regions protected from DNase I cleavage, due to ParR-strep binding, are highlighted in red boxes. (C) Foot-printing results with TraA-His protein. Compared to the non-protein control, regions where TraA-His protects the DNA from DNase I cleavage are marked with red boxes. (D) Foot-printing results with TraD-His protein. Protected regions identified by comparison with the control are highlighted, indicating where TraD interacts with the DNA. The orange lines indicate internal standards for alignment, and the blue peaks represent DNase I-cleaved DNA fragments. The areas within red boxes show regions of reduced DNase I cleavage, indicating specific binding of each respective protein compared to the "Top Non" control. The raw data were analyzed using TraceViewer software.

**Figure S4.**
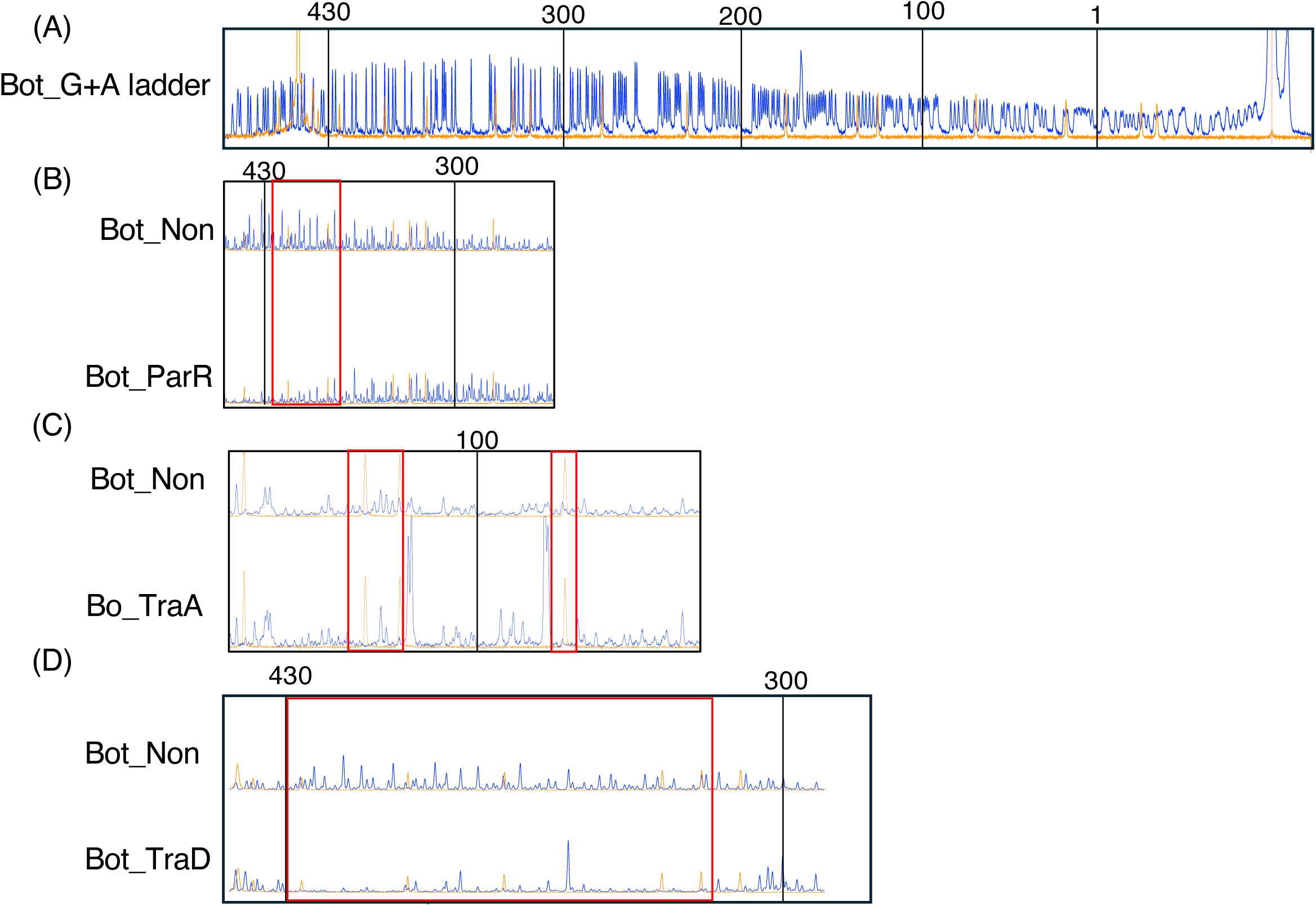
DNase I foot-printing analysis of the *oriT* region on the bottom strand. This figure illustrates the results of DNase I foot-printing assays conducted to identify protein binding regions along the bottom strand of the oriT DNA. The blue peaks represent DNA cleaved by DNase I, and the orange peaks represent internal standards for alignment. (A) The G+A ladder serves as a reference to locate specific nucleotide positions on the DNA strand, corresponding to the oriT region depicted in Fig. 5(C), enabling precise mapping of protein-DNA interactions. (B) Foot-printing results with ParR-strep protein. The non-protein control ("Bot Non") shows the baseline cleavage pattern, and regions protected from DNase I cleavage, due to ParR-strep binding, are highlighted in red boxes. (C) Foot-printing results with TraA-His protein. Compared to the non-protein control, regions where TraA-His protects the DNA from DNase I cleavage are marked with red boxes. (D) Foot-printing results with TraD-His protein. Protected regions identified by comparison with the control are highlighted, indicating where TraD-His interacts with the DNA. The orange lines indicate internal standards for alignment, and the blue peaks represent DNase I-cleaved DNA fragments. The areas within red boxes show regions of reduced DNase I cleavage, indicating specific binding of each respective protein compared to the "Bot Non" control. The raw data were analyzed using TraceViewer software.

**Figure S5.**
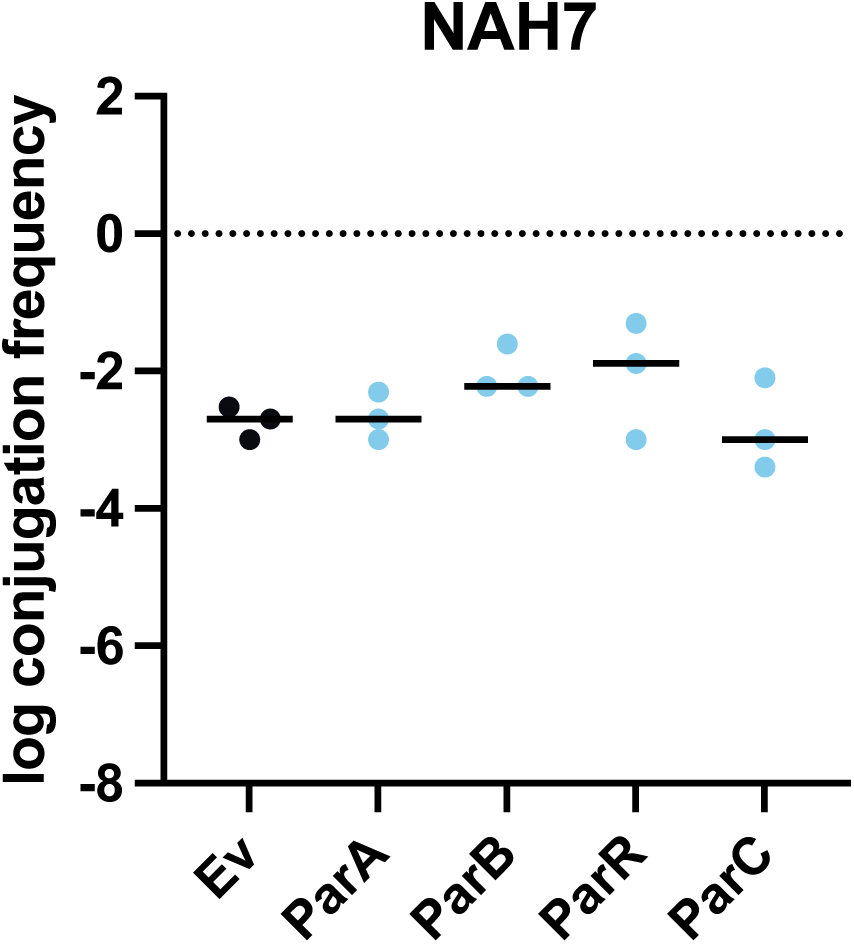
Conjugative transfer frequency of NAH7 in strains expressing Par proteins. The graph depicts the frequency of conjugative transfer of NAH7 in strains expressing different partitioning proteins (ParA, ParB, ParR, ParC) compared to an empty vector control (EV). Transfer frequency (Tc/D, number of transconjugants per donor) was measured. The data represent at least three independent biological replicates, with individual data points shown on the bar plot. The mean values are displayed along with standard deviations represented as error bars.

**Figure S6.**
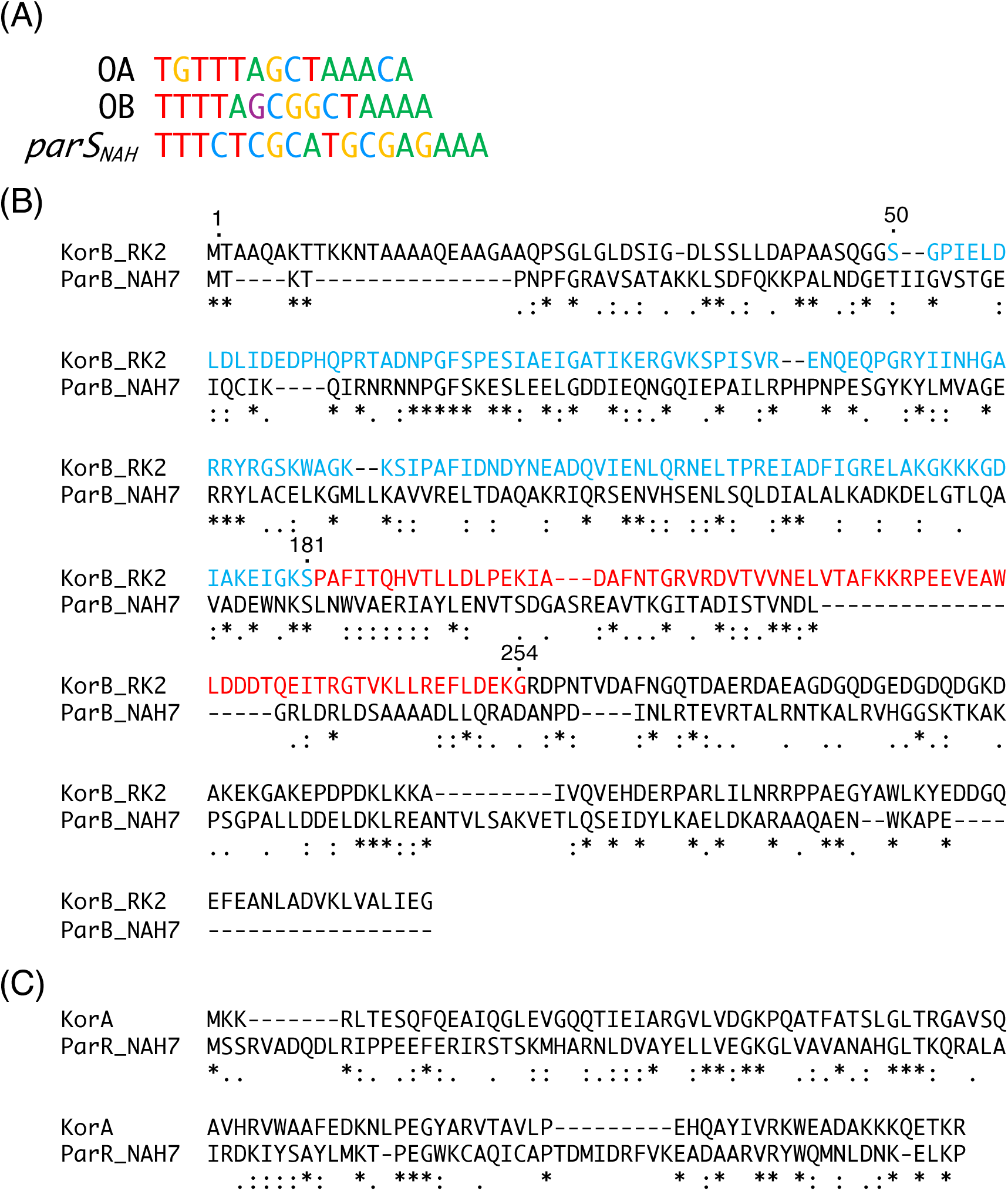
Sequence comparison of ParB_NAH7 with KorB_RP4 and ParR_NAH7 with KorA_RP4. (A) The sequences OA and OB are from KorA and KorB binding sites respectively, and *parS_NAH_* is the binding site for ParB in the NAH7 plasmid. (B) Amino acid sequence alignment of KorB from RK2 and ParB from NAH7. The alignment highlights regions of homology, with identical residues marked by asterisks (*), conserved substitutions by colons (:), and semi-conserved substitutions by periods (.). The sequences show that KorB and ParB share significant similarity, suggesting similar functions in partitioning systems. Notable domains, such as the N-terminal domain, are highlighted in blue and red, indicating DNA-binding domain. (C) Amino acid sequence alignment of KorA from RK2 and ParR from NAH7.

**Figure S7.**
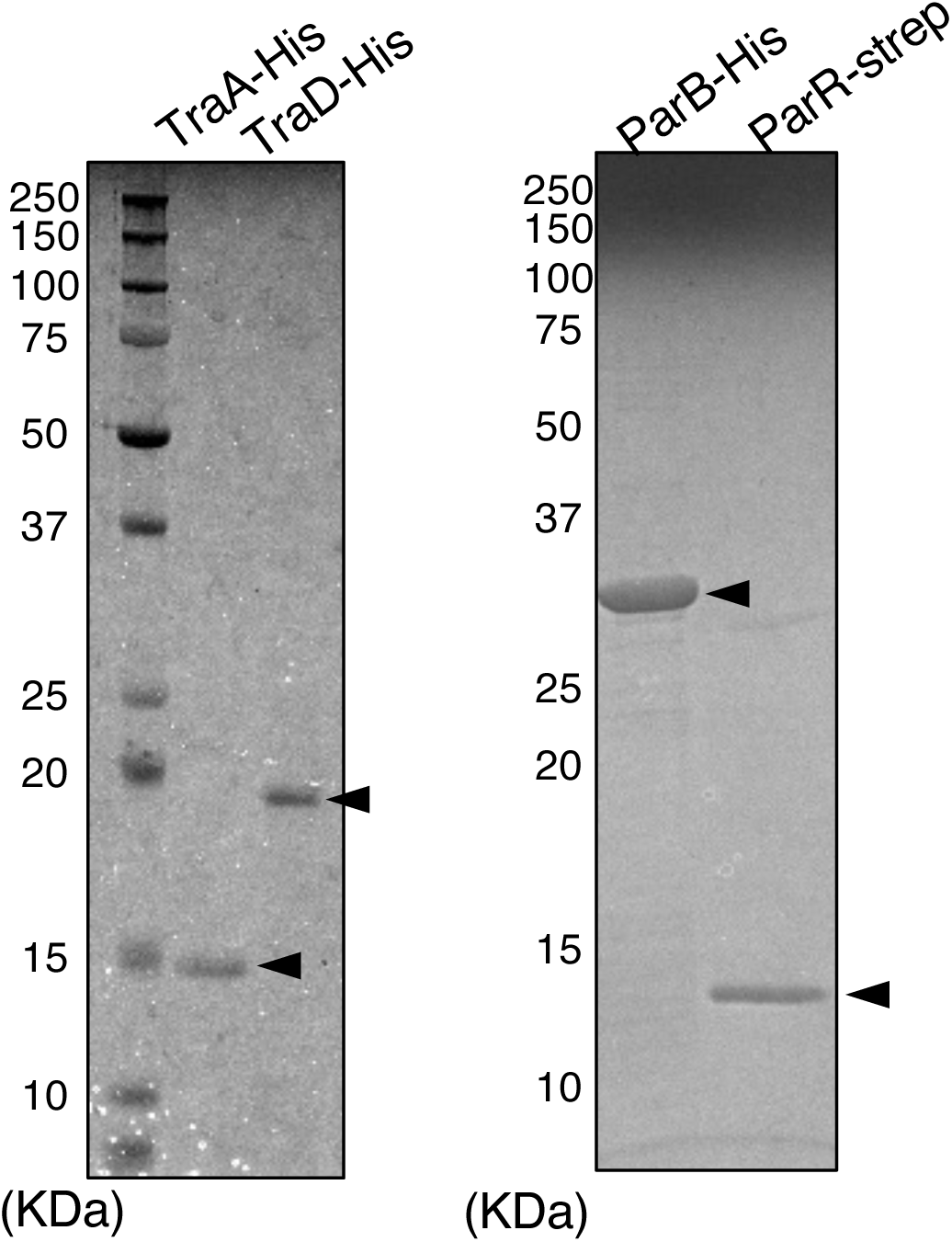
SDS-PAGE analysis of purified proteins. SDS-PAGE analysis of the purified His-tagged proteins and strep-tagged protein used in this study. The gel on the left shows the purified TraA-His and TraD-His proteins, while the gel on the right shows the purified ParB-His and ParR-Strep proteins. The molecular weight markers (in kilodaltons, kDa) are shown on the left side of each gel. Arrowheads indicate the expected bands for each purified protein.

**Table S1. Oligonucleotides used in this study.**

**Table S2. The conservation of *parABRC* within IncP-9 plasmids.**

**Table S1.**
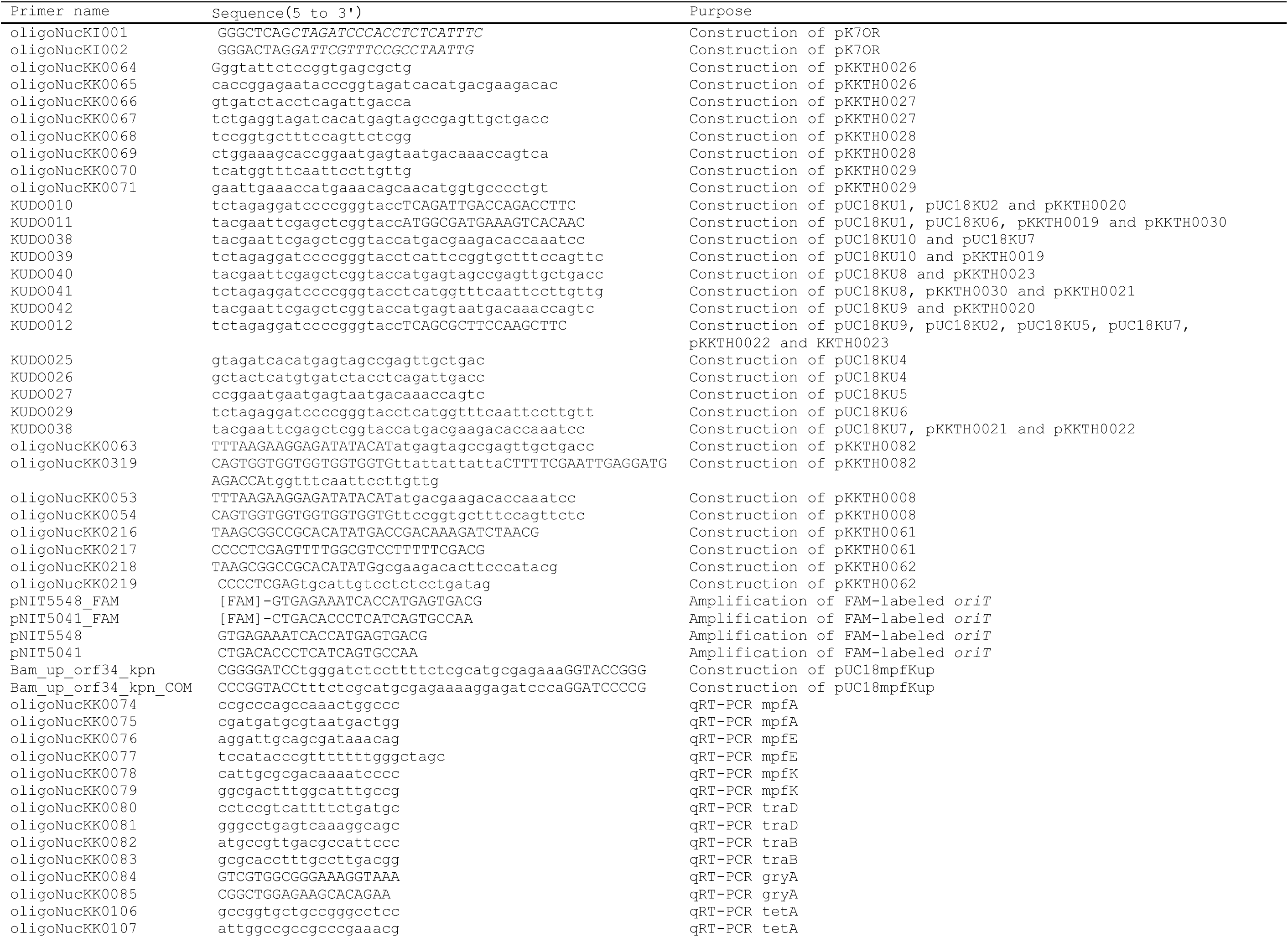
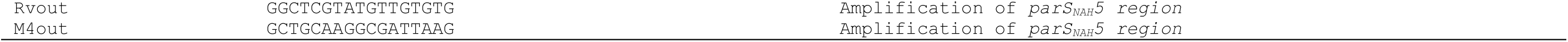
Primers used in this study^*a*^.

